# Waves of transcription drive erythroid differentiation and launch the NRF2-activated antioxidant program

**DOI:** 10.1101/2024.07.12.603281

**Authors:** Ingrid Karppi, Jenny C. Pessa, Adelina Rabenius, Samu V. Himanen, Bina Prajapati, Emilia Barkman Jonsson, Maria K. Vartiainen, Lea Sistonen, Anniina Vihervaara

**Affiliations:** Faculty of Science and Engineering, Cell Biology, Åbo Akademi University, Turku, Finland; Turku Bioscience Centre, University of Turku and Åbo Akademi University, Turku, Finland; Department of Gene Technology, KTH Royal Institute of Technology, Science for Life Laboratory, Stockholm, Sweden; Institute of Biotechnology, HiLIFE, University of Helsinki, Helsinki, Finland

**Keywords:** antioxidation, enhancer, erythroid differentiation, GABPA, GATA1, globin, glutathione, HEMGN, NRF2, PRO-seq, thioredoxin, transcription regulation

## Abstract

Transcriptional reprogramming drives differentiation and coordinates cellular responses. While mRNA expression in distinct cell types has been extensively analyzed, the mechanisms that control RNA synthesis upon lineage specifications remain unclear. Here, we induce erythroid differentiation in human cells, track transcription and its regulation at nucleotide-resolution, and identify molecular mechanisms that orchestrate gene and enhancer activity during erythroid specification. We uncover waves of transcription and reveal that a brief differentiation signal launches sustained and propagating changes in RNA synthesis and mRNA expression over cell divisions. NRF2, a strong *trans*-activator upon oxidative stress, drives erythroid differentiation without a detectable increase in reactive oxygen species. In erythroid precursors, NRF2 induces architecturally primed, differentiation-linked enhancers, and genes encoding globin and antioxidant proteins. Projecting signal-induced transcription to DNA accessibility and mRNA expression in single human bone marrow cells, reveals ordered activation of myeloid (GABPA) and erythroid (GATA1, TAL1 and HEMGN) factors in lineage-specification, followed by NRF2-triggered antioxidant response in the late erythroid cells. This study establishes molecular mechanisms that prime, execute, and temporally coordinate RNA synthesis during erythroid differentiation. Furthermore, we show that master regulators of differentiation and stress co-orchestrate erythropoiesis and produce the antioxidant machinery before erythroid cells mature to oxygen transporting enucleated erythrocytes.

## Introduction

Transcription from protein-coding genes, non-coding genes and enhancers is carefully regulated to establish cellular identities and adjust to internal and external conditions^1^. Differentiation of stem cells is vital for development, tissue repair, and turnover of cells throughout life. Human erythrocytes live approximately 120 days and are constantly replenished by division and specialization of hematopoietic stem cells (HSCs) in the bone marrow. In adults, erythropoiesis is driven by lineage-specific transcription factors GATA-binding factor 1 (GATA1), GATA2, and T-cell acute lymphocytic leukemia protein 1 (TAL1)^2–3^. The erythroid differentiation triggers activation of hemoglobin synthesis, reduction in cell size, and loss of all organelles. A robust model for erythropoiesis in human is K562 cell line, a multipotent hematopoietic progenitor derived from a patient with chronic myeloid leukemia^4–5^. K562 cells can be stimulated toward distinct myeloid lineages, including erythroid and megakaryocytic paths^6–7^. The differentiation toward the erythroid lineage is induced with hemin, a Fe^3+^-containing form of heme that binds oxygen in hemoglobin. Indeed, both heme and hemin activate erythroid differentiation *in vitro* and *in vivo*, manifested by increased hemoglobin production^5^.

Transcription factors are classified into functional categories based on their expression and inducibility^8^. The lineage-specific factors are exclusively expressed in certain cell types and direct differentiation programs with long-term changes in the chromatin state and transcription. In mammals, lineage-specific transcription factors bind predominantly to enhancers, characterized by open chromatin, H3K4me1 and H3K27ac histone modifications, and synthesis of enhancer RNA (eRNA)^9–10^. In comparison, transcriptional regulators of stress responses are ubiquitously expressed and exhibit low basal activity. These potent *trans*-activators are rapidly induced upon adverse conditions and can reprogram transcription within minutes^11^. Proteotoxic stress activates heat shock factor 1 (HSF1) to provoke instant, pervasive, and transient transcriptional reprogramming, causing rapid increase in chaperone expression^12–16^. High levels of reactive oxygen species (ROS) release nuclear factor erythroid 2 (NFE2)-related factor 2 (NRF2, *NFE2L2*) from kelch-like ECH-associated protein 1 (KEAP1). The release from KEAP1 allows NRF2 to evade degradation and launch the antioxidant program^17–18^. Both stress and differentiation incite drastic changes to RNA and protein expression and require careful regulation of homeostasis^19^. To date, the interplay between lineage-specific and stress-induced transcription factors is poorly understood.

To uncover mechanisms and kinetics of transcriptional reprogramming during differentiation, we induced K562 cells toward the erythroid lineage and tracked RNA Polymerase II (Pol II) progression across genes and enhancers at nucleotide-resolution using Precision Run-On sequencing (PRO-seq)^20^. Hemin-induced erythroid differentiation proceeded in waves of transcription, coordinated predominantly at the rate-limiting step of initiation. A brief hemin signal launched instant and late waves of transcription that continued propagating in the daughter cells, even days after the hemin removal, demonstrating continued differentiation in the absence of the initial trigger. The transcriptional responsiveness to erythroid differentiation was primed *via* promoter-specific and enhancer-specific chromatin architectures, involving lineage-specific factors, signal-induced activators, and chromatin remodelers. To our surprise, erythroid differentiation activated NRF2 without a detectable increase in ROS. Specifically, NRF2 potentiated globin expression, launched the antioxidant program, and activated architecturally primed enhancers that contained NRF2 motifs upstream the +1 nucleotide. Projecting the chromatin architecture, signal-induced transcription kinetics and mRNA expression to single cell RNA-seq and ATAC-seq (SHARE-seq) data^21^ in human bone marrow, revealed ordered activation of myeloid (GA-binding protein subunit alpha, GABPA), and erythroid (GATA1, TAL1 and hemogen, HEMGN) factors to drive lineage-specification, followed by NRF2-triggered production of antioxidant machinery in the late erythroid cells. This study identifies the molecular logic that coordinates transcription during human erythropoiesis. Moreover, master regulators of differentiation and stress are shown to co-orchestrate transcriptional reprogramming during erythroid specification and produce the antioxidation machinery in the late erythroid cells before the cells mature to oxygen transporting erythrocytes that are enucleated and exposed to ROS.

## Results

### Transcriptional waves of differentiation

To dissect mechanisms that reprogram transcription during lineage specification, we activated erythroid differentiation in K562 cells with hemin. As expected, hemin turned K562 cells dark red (Fig. 1A), reflecting differentiation-induced production of hemoglobin. Transcriptional reprogramming was measured with PRO-seq that labels active sites of transcription with biotinylated nucleotides and reports Pol II progression across genes and enhancers at nucleotide-resolution^20^. To capture both instant and sustained changes in RNA synthesis, we performed PRO-seq during a comprehensive time course of differentiation, ranging from early (15-60 min) to late (24 h) hemin treatments, and including a 48-h recovery from either a short (60 min) or a long (24 h) treatment (Fig. 1B). Each sample was prepared in two biological replicates, which provided highly similar Pol II densities across the genes (rho>0.98, Fig. S1A), including *HSPA1A* and *HSPA1B* (Fig. S1B-C) that were previously shown to be hemin-induced^22^.

**Figure 1.**
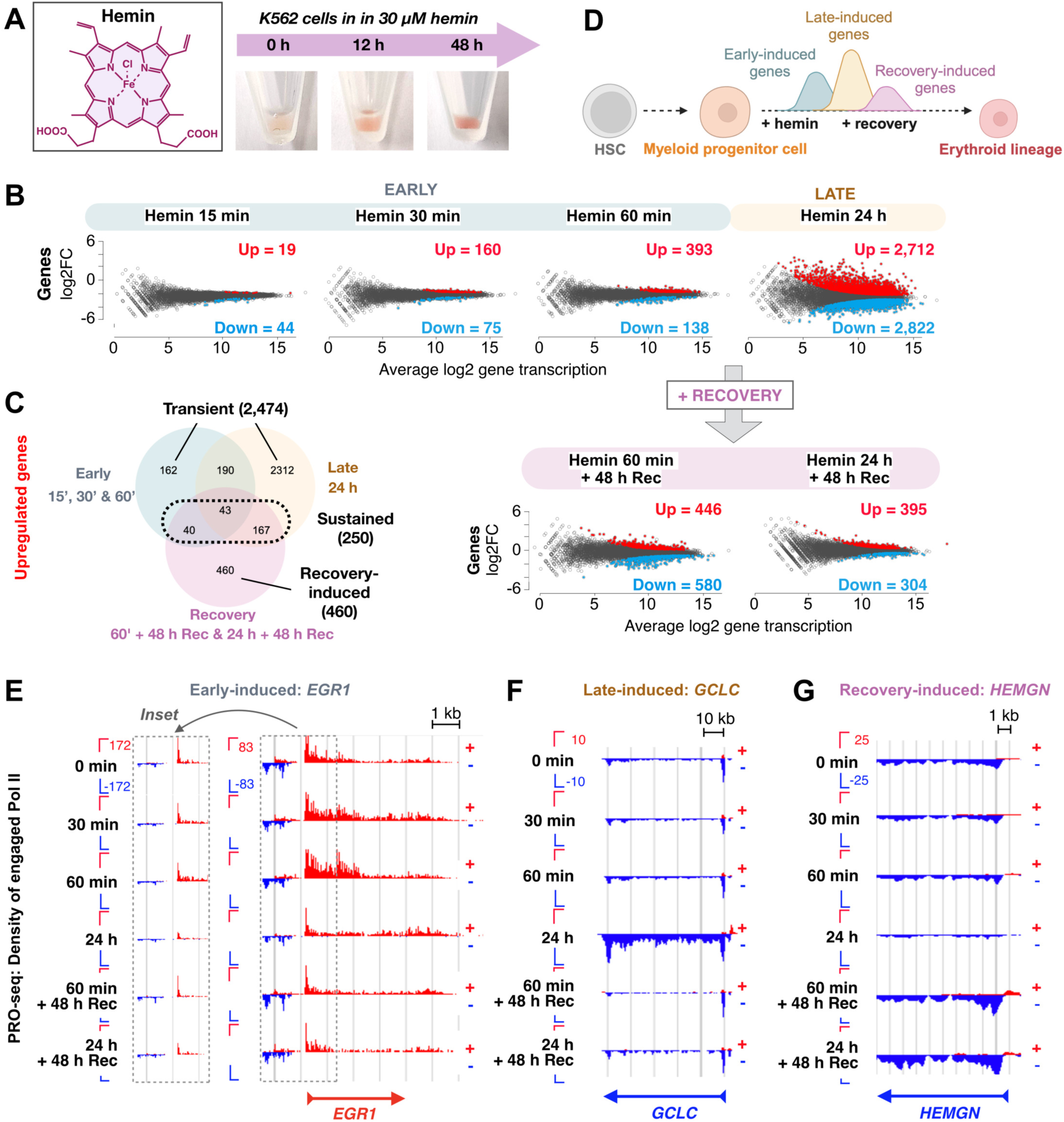
Hemin-induced differentiation launches waves of nascent transcription. **A)** Structure of hemin (left) and photographs of hemin-treated K562 cells (right) showing the darkening red color characteristic for increased globin production. **B)** Genes with significantly (p<0.01) induced (red) or repressed (blue) nascent transcription upon hemin-induced differentiation as identified with DESeq2 and compared to untreated K562 cells. **C)** Overlap or early-, late-, and recovery-induced genes. **D)** Schematic presentation of the distinct transcriptional waves upon erythroid differentiation. **E-G)** Genome browser examples of hemin-induced genes. **E)** Early growth response protein 1 (*EGR1*) is induced during the first hour of hemin-treatment, while **F)** glutamate-cysteine ligase catalytic subunit (*GCLC*) is induced upon 24 h, and **G)** hemogen (*HEMGN*) after 48-h recovery. Red tracks denote RNA synthesis on the plus strand, blue tracks on the minus strand. The inset in **E** shows divergent transcription and promoter-proximal Pol II pausing at *EGR1*. Related Supplementary Figure 1.

Inducing erythroid differentiation launched extensive transcriptional reprogramming of genes (Fig. 1) and enhancers (Fig. 2A-C). Genes with statistically significant change in RNA synthesis (hereafter induced and repressed) were detected already at 15 min of hemin treatment, and the transcriptional changes gradually propagated to over 5,000 genes at 24 h (Fig. 1B, Dataset 1). Many genes were only transiently induced or repressed, demonstrating that erythroid differentiation proceeded in transcriptional waves (Figs. 1C-D and S2A). An example of an instantly but transiently activated gene (Fig. 1E) encodes the early growth response protein 1 (EGR1), a transcription factor that drives rapid changes in cells states, including in differentiation^23–25^. One of the genes activated during the late wave encodes the catalytic subunit of glutamate-cysteine ligase (GCLC) (Fig. 1F). GCLC is the rate-limiting enzyme for synthesis of glutathione (GSH), an essential component of the cellular redox system^26^. Gene ontology (GO) analyses (Dataset 2) uncovered the early transcriptional wave to be enriched for erythrocyte development (fold enrichment, fe, 7.49; false discovery rate, FDR, 2.94E-02) and the late wave for autophagy (fe 2.99; FDR 5.07E-18), particularly mitophagy (fe 4.46; FDR 1.28E-02). This suggested that erythroid properties were induced by hemin, and that in the late time point, cells activated a program mimicking the terminal stages of erythropoiesis where autophagy contributes to organelle removal^27^.

**Figure 2.**
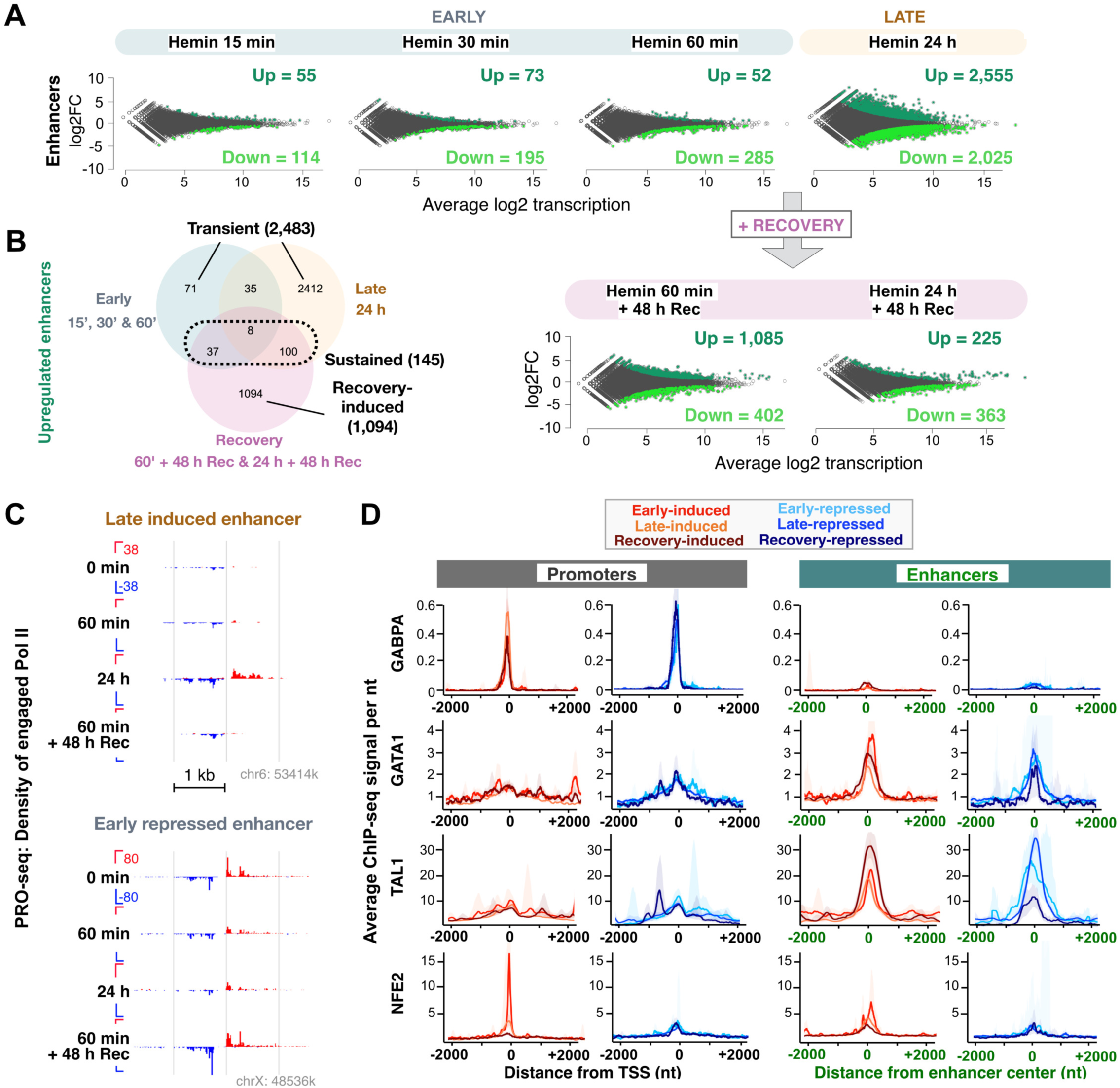
Erythroid differentiation triggers rapid and long-lasting changes in eRNA synthesis. **A)** MA-plots showing enhancers with significantly (p<0.01) induced (dark green) or repressed (light green) eRNA synthesis upon hemin-induced differentiation, as identified with DESeq2 and compared to untreated K562 cells. **B)** Overlap between early-, late-, and recovery-induced enhancers. **C)** Genome browser examples of late-induced (top) and early-repressed (bottom) enhancers. **D)** Average ChIP-seq intensities of the indicated lineage-specific transcription factors in untreated K562 cells. The average intensities are shown for early- (red), late- (orange) and recovery- (mahogany) induced, as well as early- (light blue), late- (blue) and recovery- (dark blue) repressed promoters and enhancers. The shaded area around the average denotes 12.5-87.5% confidence interval. The ChIP-seq data is from the ENCODE^36^. Related Supplementary Figures 2 and 3.

### A brief trigger seeds persistent changes

The hemin-induced differentiation in K526 cells has been considered reversible, as hemoglobin levels gradually diminish after hemin removal^28^. However, tracking transcription after removing the hemin (hereafter termed recovery), showed that a remarkable part of the changes in nascent RNA synthesis was not restored (Fig. 1B). On the contrary, transcriptional changes triggered by the hemin continued in the absence of the signal, as more genes were induced and repressed after a 48-h recovery than at 60-min hemin treatment (Fig. 1B). In log phase, K562 cells divide every 22-26 h^16,28^, suggesting that the differentiation-induced transcriptional changes propagated over cell divisions. We confirmed the earlier findings^28–29^ that a brief hemin treatment neither interfered with the cell cycle progression nor increased apoptosis (Fig. S2B). Instead, K562 cells continued dividing through the recovery (Fig. S2C), showing that hemin-induced transcriptional changes propagated over cell divisions.

A notable number of genes were significantly induced or repressed only after the recovery (Figs. 1C and S2A), denoting a separate transcriptional wave in the daughter cells (termed recovery-induced wave). The recovery-induced genes were enriched for erythrocyte differentiation (fe 3.92; FDR 1.83E-02, Dataset 2), indicating that the cells continued in the differentiation path even in the absence of the signal that triggered the change. One of the genes whose transcription was induced during the recovery encodes HEMGN (Fig. 1G) that promotes erythroid maturation and heme biosynthesis^30–31^. These findings imply that even a brief signal of erythroid differentiation can, through a series of transcriptional events, direct cells toward a new identity (Fig. 1D).

### Rewiring enhancer transcription

Enhancers are key regulators of gene expression during lineage specification, and eRNA synthesis correlates with enhancer activity^9–10^. We scanned our PRO-seq data for the characteristic pattern of eRNA synthesis^32^ and quantified transcription at enhancers during hemin-induced differentiation. Erythroid differentiation triggered profound reprogramming of eRNA synthesis that, similarly to gene transcription, followed propagating waves of induction and repression (Figs. 2A-C, S2D, Dataset 1). Only 52 enhancers were induced at 60 min of hemin treatment, but after the subsequent 48-h recovery, more than a thousand enhancers increased eRNA synthesis (Figs. 2A-B, S2D), which indicated that a transient signal of differentiation triggered long-lasting changes in gene and enhancer transcription. eRNA synthesis has been detected to precede or co-occur with the target gene induction depending on the stimulation^33–34^. During erythroid differentiation, the transcriptional activation and inactivation of genes and enhancers followed a similar temporal pattern, exemplified by *GCLC* (Fig. 1F) and its nearest enhancer (Fig. 2C, upper panel). Moreover, both early-responding genes and enhancers maintained the course of transcriptional change throughout the differentiation, while most late-responding genes and enhancers returned to their basal transcription during the recovery (Fig. S2E).

### Priming promoters and enhancers

Dynamic reprogramming of transcription depends on chromatin accessibility, transcription factor binding, and the state of the transcription machinery^8,35^. To assess how cells elicit transcriptional responses upon signal-induced differentiation, we investigated chromatin architecture at hemin-responsive promoters and enhancers using the vast ENCODE data in K562 cells^36^. Promoters, and to a lower extent enhancers, were in an open chromatin state prior to the hemin treatment (Fig. S3A). Moreover, differentiation-activated promoters and enhancers were flanked by high levels of H3K4me1 (Fig. S3B), a chromatin mark associated with a poised state at both regulatory elements^37–38^. In comparison, H3K4me3 was considerably higher at promoters than enhancers (Fig. S3B).

Next, we assessed the binding of lineage-specific factors for myeloid or erythroid differentiation and found their strong preferences for either promoters or enhancers (Fig. 2D). GABPA, an essential factor for differentiation into the myeloid lineage^39^, was abundant across promoters, but almost absent from enhancers (Fig. 2D). On the contrary, GATA1 and TAL1 occupied primarily enhancers (Fig. 2D). Erythroid-specific transcription factor NFE2 was enriched at early-induced promoters, detectable at early-induced enhancers, and virtually absent from the repressed regulatory elements (Fig. 2D). Beyond the lineage-specific factors, activating transcription factor 3 (ATF3), shown to bookmark genes for rapid activation^40^, occupied early-induced promoters (Fig. S3C). Small Maf transcription factors, including MAFF, that can switch from a repressor to an activator upon interacting with NRF2^41^, were abundant at the induced enhancers (Fig. S3D). To further highlight the distinct chromatin architectures at differentiation-responsive promoters and enhancers, several chromatin remodelers preferred either enhancers or promoters, as demonstrated with enhancer-specific enrichment of chromodomain helicase 1 (CHD1) and promoter-specific occupancy of CHD2 (Fig. S3E). These results identified that differentiation-linked promoters and enhancers resided in accessible and poised chromatin state prior to the hemin treatment. However, promoters and enhancers were primed for signal-induced erythroid differentiation by distinct chromatin remodelers and lineage-specific transcription factors.

### The key rate-limiting step is initiation

Progression of Pol II through genes and enhancers is regulated at the rate-limiting steps of transcription^35^. Of these steps, transcription initiation and Pol II pause-release have emerged as the main stages at which transcriptional reprogramming across mammalian genomes is coordinated. To dissect the molecular mechanisms that control RNA synthesis during erythroid differentiation, we first quantified Pol II pause complexes at the promoter-proximal regions and along the short enhancers (Fig. S4A). Both promoters and enhancers that responded early contained abundantly of Pol II before the stimulation (Fig. S4A), which together with the accessible chromatin (Fig. S3A) and the lineage-specific factors (Fig. 2D), demonstrates architectural priming of the rapidly responding regulatory elements.

Next, we monitored Pol II progression from the initiation, through the promoter-proximal pause region, into the productive elongation (Fig. 3A-B). At repressed genes, the average Pol II density decreased from the initiation to the end of the gene, showing reduced Pol II pausing as the differentiation proceeded (Fig. 3A-B). This decrease in Pol II density along the whole gene evidenced diminished entry of the transcription machinery into the gene and established that the main rate-limiting step for transcriptional repression upon erythroid differentiation was initiation. At the induced genes, Pol II density increased downstream of the pause, but the average pause density remained the same, both at early- (Fig. 3A) and late- (Fig. 3B) activated genes. This increased entry of Pol II into productive elongation can be accomplished by either a higher rate of initiation coupled to efficient Pol II pause-release, or reduced pre- mature termination at the pause, which enables a higher fraction of the paused Pol II to enter productive elongation.

**Figure 3.**
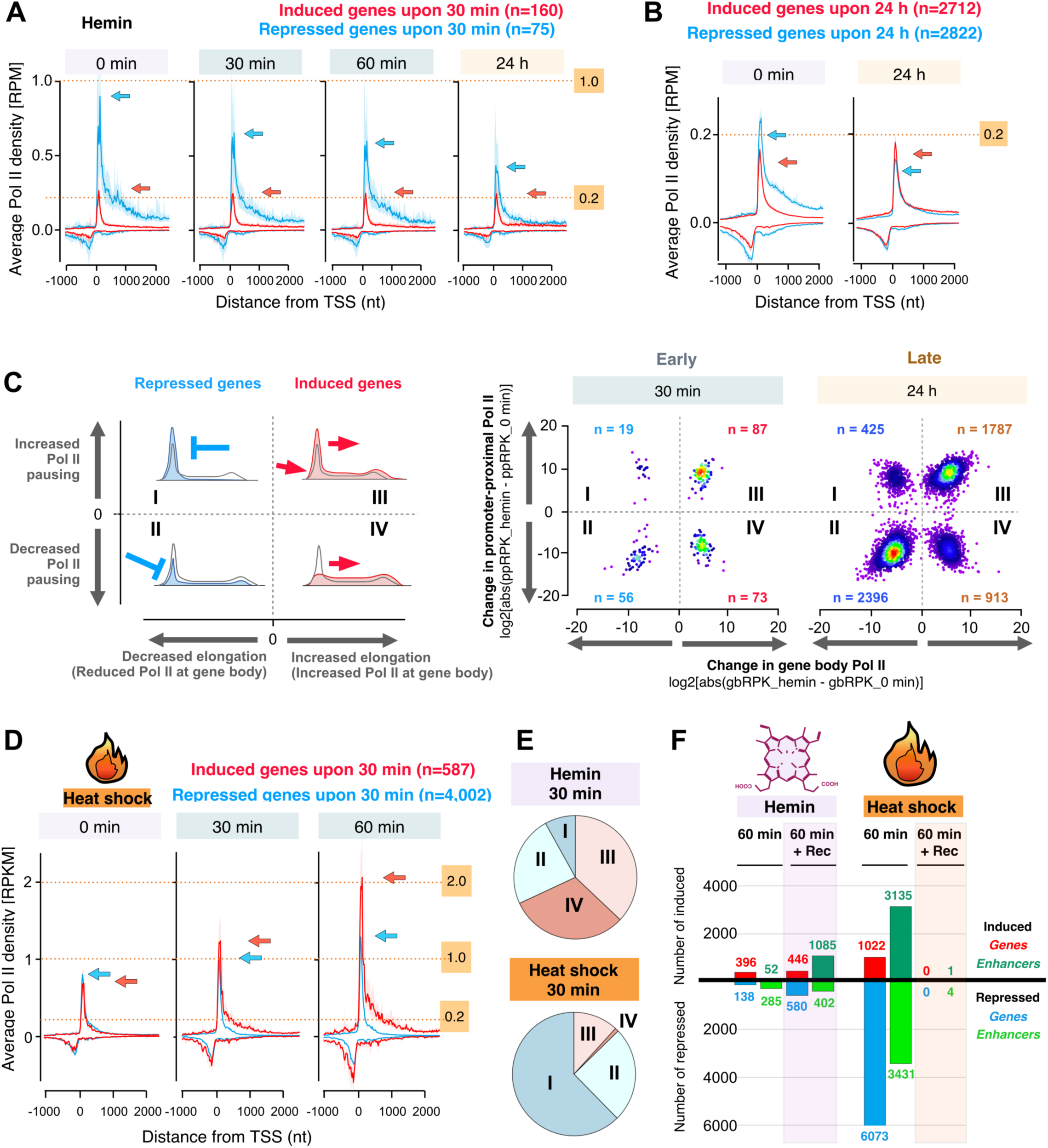
Differentiation-induced transcriptional reprogramming is controlled at initiation. **A-B)** Average intensity of transcriptionally engaged Pol II at promoter-proximal regions of induced (red) and repressed (blue) genes. The distinct panels follow the change in Pol II engagement during hemin-induced differentiation. Blue arrow: The highest Pol II pause signal at hemin-repressed genes in each time point. Red arrow: The highest Pol II pause-signal at hemin-induced genes in each time point. **C)** Schematic key (left) of the scatter plots (right) deciphering the change in Pol II occupancy at the promoter-proximal pause region and the gene body. Each gene is indicated with a sphere and positioned based on the change in Pol II pausing (y-axis) and productive elongation (x-axis). Genes in the top left square (I) are repressed by inhibiting the release of Pol II into productive elongation (increased Pol II pausing, reduced elongation). In the bottom left square (II), genes are repressed by decreased transcription initiation (less Pol II at the pause and along the gene). Genes in the top right square (III) are induced by a higher rate of initiation (increased Pol II at the pause and along the gene). In the bottom right square (IV), the rate of pause-release increases more than the rate of initiation (reduced pausing, increased elongation). **D)** Average intensity of transcriptionally engaged Pol II at promoter-proximal regions of heat-induced (red) and heat-repressed (blue) genes. The arrows indicate the highest Pol II pausing at heat-repressed (blue) and heat-induced (red) genes. **E)** Comparison of changes in the rates of initiation and pause-release at differentially transcribed genes upon erythroid differentiation (upper panel) and heat shock (lower panel). The roman numerals (I-IV) correspond to the groups depicted in **C**. **F)** Transcriptional reprogramming of genes and enhancers compared during erythroid differentiation (hemin) and acute stress (heat shock) in K562 cells. The number of genes and enhancers that remain significantly changed after a-48h recovery from the respective treatment are shaded. The PRO-seq data in heat shocked K562 cells originates from reference^16^. Related Supplementary Figure 4.

To assess how Pol II is controlled at individual genes, we quantified changes in the rates of initiation and pause-release at each repressed (groups I and II) or induced (groups III and IV) gene (Fig. 3C). We found that 75% of the early-repressed and 85% of the late-repressed genes were inhibited by decreased rate of initiation (group II, Fig. 3C). Only a few genes were repressed *via* inhibition of the pause-release (group I, Fig. 3C), shown as increased Pol II density at the pause. At the induced genes, 55% of the early and 66% of the late responders increased the rates of initiation and pause-release (group III), elevating Pol II engagement across the gene (Fig. 3C). However, at a considerable fraction of the induced genes, the rate of pause-release increased more than the rate of initiation (group IV), which reduced Pol II pausing. Taken together, initiation emerged as the main step that coordinated transcriptional reprogramming upon erythroid differentiation. Additionally, the rate of pause-release increased at activated genes, enabling efficient progression of the transcription machinery through the pause-region into productive elongation.

### Mechanisms of persistent reprogramming

The initiation-driven transcriptional reprogramming during erythroid differentiation (Figs. 1-3) was profoundly different from the instant and transient transcriptional changes reported upon acute stress^11,16,18^ and during macrophage efferocytosis^42^. To directly compare the mechanisms of transcriptional reprogramming upon acute stress *versus* differentiation, we generated Pol II density profiles (Fig. 3D) and measured initiation and pause-release (Figs. 3E, S4B-C) in K562 cells exposed to heat shock. Unlike erythroid differentiation, heat shock instantly repressed thousands of genes by inhibiting the Pol II pause-release, which accumulated Pol II at the pause (Fig. 3D). Gene-by-gene quantification showed that 71% of the heat-repressed genes reduced the rate of pause-release (group I, Figs. 3E and S4B-C). Instead, the heat-induced genes displayed high Pol II pausing prior to stress (Fig. 3D) and a massive increase in initiation and pause-release upon activation, which was manifested by 92% of heat-induced genes increasing both initiation and pause-release (group III, Figs. 3E and S4B-C). The contrast between the mechanisms that coordinated Pol II molecules upon differentiation *versus* stress was most profound in the recovery phase (Fig. 3F). While hemin launched slowly propagating but persistent transcriptional changes over cell divisions (Fig. 3F, left side), the heat-induced reprogramming was rapid and transient (Fig. 3F, right side). These results imply drastically different strategies for transcriptional reprogramming depending on the stimulation. Transcriptional stress responses are instant, transient, and coordinated at the Pol II pause-release, whereas erythroid differentiation follows slow kinetics, commits cells to long-lasting transcriptional changes, and redistributes Pol II across the genome by coordinating transcription initiation.

### NRF2 drives erythroid differentiation

Erythropoiesis is regulated by networks of lineage-specific transcription factors^2^. To identify transcription regulators activating the distinct waves, we performed transcription factor prediction analyses. As expected, the hemin-responding genes were significantly enriched with many known targets of GATA1, GATA2 and GABPA (Fig. 4A, Dataset 3). To our surprise, Enrichr also identified NRF2, the *trans*-activator of the antioxidant program, as a putative transcription factor driving erythroid differentiation (Fig. 4A, Dataset 3). Tens of NRF2 target genes were activated upon the early, and hundreds upon the late, wave of transcription (Fig. 4A). These genes included the widely studied NRF2 targets^41^ essential for the three main antioxidant pathways: 1) synthesis and metabolism of the antioxidant GSH, 2) thioredoxin (TXN)-mediated antioxidation, and 3) NADPH regeneration (Fig. 4B-C). Some of the NRF2 target genes reacted instantly, including solute carrier family 7 member 11 (*SLC7A11*) (Fig. 4D). *SLC7A11* encodes antiporter xCT that imports cysteine, an essential component in GSH^43^. Overall, we found substantial induction of NRF2 target genes that are central for the GSH pathway, such as *GCLC* (Fig. 1F), glutamate-cysteine ligase modifier subunit (*GCLM*), and glutathione-disulfide reductase (*GSR*) (Figs. 4B-C and S5A, Dataset 1). Of NRF2 target genes, also *TXN* (Fig. S5B) and its reductases thioredoxin reductase 1 (*TXNRD1*) and sulfiredoxin 1 (*SRXN1*) were activated (Fig. 4B and Dataset 1). GSH and TXN are antioxidants that donate electrons to oxygen radicals. This ROS neutralization converts GSH to its oxidized and dimerized GSSH form, and TXN to TXN-S_2_ (Fig. 4C). To enable a new round of ROS neutralization, GSSH and TXN-S_2_ need to be reduced back to GSH and TXN in a process that requires NADPH^44^ (Fig. 4C). In accordance, several NRF2 targets that encode enzymes involved in NADPH production were activated (Fig. 4B-C, Datasets 1 and 3), completing the cycle for ROS neutralization by GSH and TXN.

**Figure 4.**
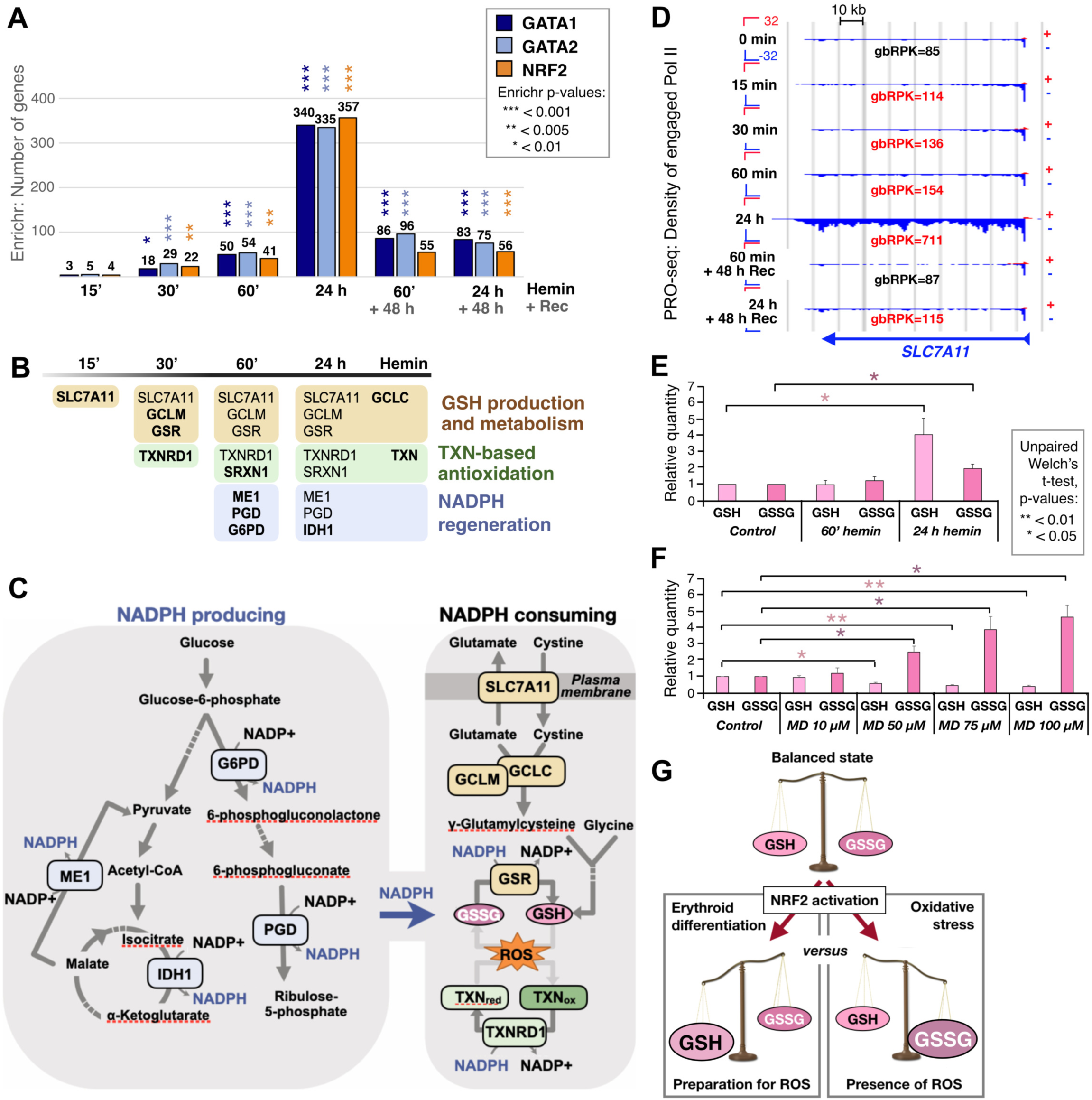
NRF2 target genes are induced upon erythroid differentiation. **A)** Number of GATA-binding factor 1 (GATA1), GATA2, and nuclear factor erythroid 2-related factor 2 (NRF2) targets among induced and repressed genes, as detected with Enrichr. Significant p-values from Enrichr analysis are indicted. **B)** Time of NRF2 target gene induction upon erythroid differentiation. The colors indicate the three main pathways of antioxidant response: glutathione (GSH) production and metabolisms (light brown), thioredoxin (TXN)-based antioxidation (green) and NADPH production (blue). The bolded text indicates the earliest time point the gene’s nascent transcription is significantly induced. **C)** Schematic representation of the three main antioxidant pathways. Intermediates (bolded text) and hemin-induced NRF2 targets (rounded rectangles) are indicated. Dashed lines indicate intermediates not shown. **D)** Genome browser illustration of the induction of NRF2 target gene solute carrier family 7 member 11 (*SLC7A11*) upon hemin-induced erythroid differentiation. RNA synthesis as PRO-seq reads per kilobase of gene body (gbRPK) are shown for each time point, indicating significant induction in red. **E)** Relative quantity of GSH and GSSG in K562 cells treated with 30 μM hemin for 60 min or 24 h. n=3. *p<0.05. **F)** Relative quantity of GSH and GSSG in K562 cells treated with the oxidative stress inducer menadione (MD) for 60 min. n=3. *p<0.05, **p<0.01. **G)** Schematic illustration comparing the ratio of protective GSH per oxidized GSSG (GSH/GSSG) in erythroid differentiation (left) *versus* oxidative stress (right). Related Supplementary Figures 5 and 6.

NRF2 activation could indicate oxidative stress in hemin-treated K562 cells. To address whether hemin caused oxidative stress, we measured the levels of reduced GSH and oxidized GSSH upon hemin-induced erythroid differentiation (Fig. 4E) and oxidative stress provoked by menadione (Fig. 4F). Hemin-induced erythroid differentiation did not change the GSH-to-GSSG balance at 1 h (Fig. 4E). Instead, hemin treatment significantly elevated the levels of the protective GSH at 24 h (Fig. 4E), confirming the hemin-induced production of GSH and demonstrating that erythroid differentiation activates synthesis of antioxidants without a detectable increase in the oxidative stress. In stark contrast, the strong ROS-inducing agent menadione caused a concentration-dependent increase in GSSG and decrease in GSH upon a short 1-h treatment (Fig. 4F), demonstrating ROS neutralization by GSH-to-GSSG conversion. Since mature erythrocytes are highly dependent on GSH^45^, we propose that NRF2 is activated upon erythroid differentiation in a ROS-independent manner to produce the antioxidation machinery and protect the mature, enucleated oxygen carriers against ROS (Fig. 4G, left path).

### NRF2 activates promoters and enhancers

NRF2 activity has been investigated primarily through the expression of its target genes. However, MEME-ChIP analyses identified NRF2 motifs (also knowns as AREs) at transcriptionally activated enhancers (Fig. 5A, Dataset 4). The NRF2 motif was prevalent at both sides of the enhancer center, whereas the GATA motif was evenly distributed (Fig. 5B). These sites of NRF2 motif enrichments overlap with the core initiation regions that position Pol II and orient transcription toward the sense and anti-sense directions^15,46^. Indeed, early- and late-induced enhancers were enriched for the NRF2 motif as compared to enhancers with unchanged eRNA expression (Fig. 5C). The prominent activation of NRF2 motif-containing enhancers revealed a large set of putative NRF2 targets and suggested that NRF2 drives transcriptional activation of differentiation-linked genes and enhancers.

**Figure 5.**
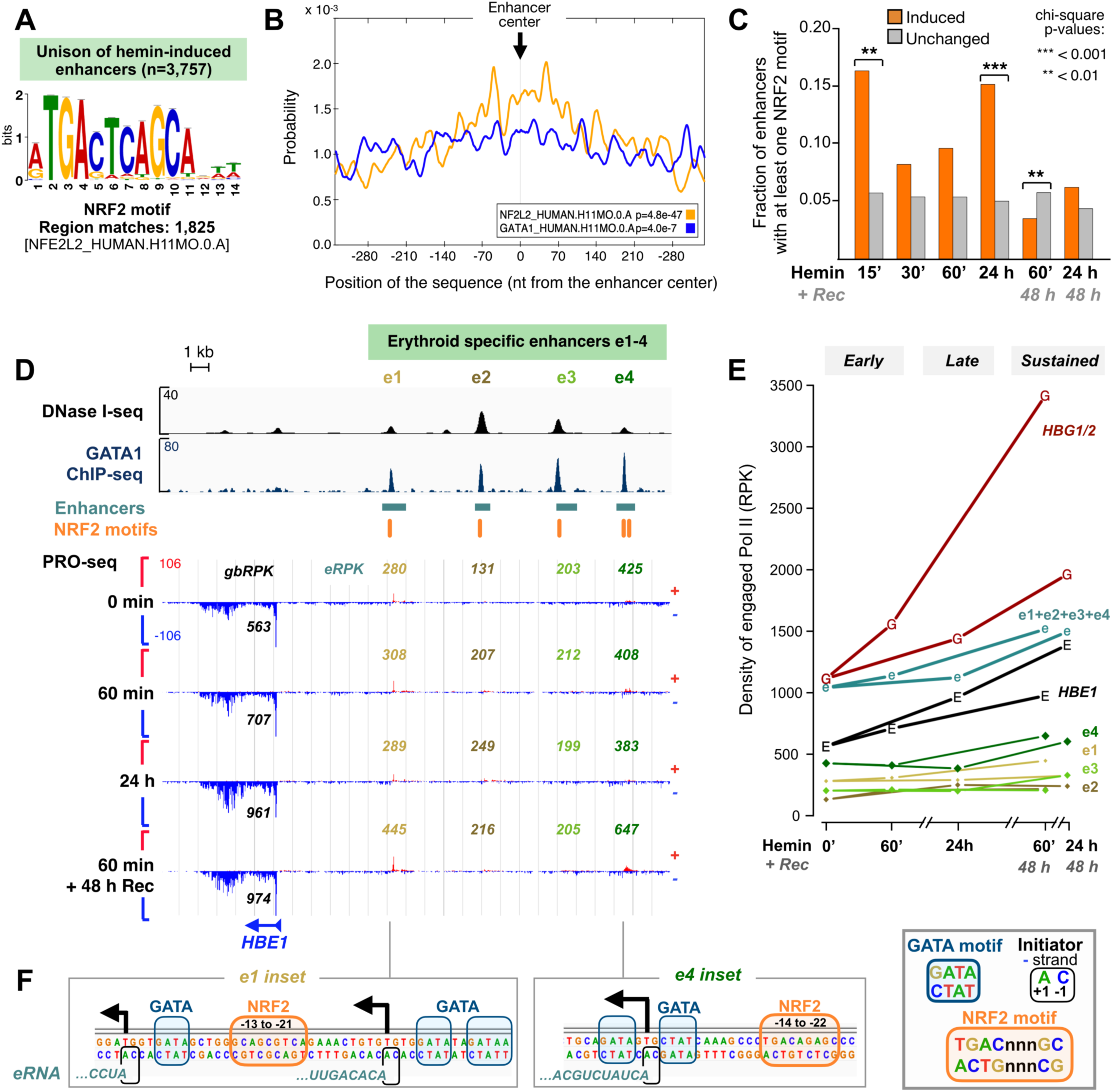
NRF2 directs erythroid differentiation *via* enhancers. **A)** The NRF2-recognized DNA motif as identified by MEME-ChIP from the unison of all hemin-induced enhancers (n= 3,757). **B)** CentriMo analysis of the probability for NRF2 and GATA1 motifs with respect to the enhancer center. **C)** Fraction of hemin-induced and unchanged enhancers that contain at least one NRF2 motif. The statistical analyses were performed with chi-square. **p<0.01, ***p<0.001. **D)** DNase I-seq, GATA1 ChIP-seq, transcribed enhancers, NRF2 motifs, and nascent transcription (PRO-seq) at the β-globin locus. The hypersensitive sites 1-4 (HS1-4), here denoted e1-4, indicate the erythroid-specific enhancers in the β-globin locus control region (LCR). The transcriptional profile of e1-4 and the nearest globin gene (*HBE1*) is shown upon 0 min, 60 min and 24 h hemin treatment, and after 48 h recovery from the 60-min hemin treatment. Quantification of *HBE1* (gbRPK; in black), and e1-4 transcription (eRPK; gold, brown, light green, dark green) are shown. **E)** Quantification of *HBE1* (black), *HBG1/2* (dark red), and e1-4 transcription (gold, brown, light green, and dark green, respectively) during erythroid differentiation. e1+e2+e3+e4 indicates the total transcription (sum eRPK) across enhancers e1-4. **F)** Insets of e1 (left) and e4 (right) from panel **D**, zooming into the GATA1 ChIP-seq summit points. Underlying architecture of transcription initiation (arrows), initiator motif, 5’-ends of eRNAs, GATA motifs and NRF2 motifs are indicated. The precise transcription start nucleotides (+1 nt) are identified from the 5’-ends of nascent transcripts shown in Figure S5C. Related Supplementary Figure 5.

To understand the functional relevance of NRF2 at enhancers, we investigated the NRF2 motif occurrence at differentiation-linked enhancers, focusing on the locus control region (LCR) of the β-globin gene cluster. The LCR contains four erythroid-specific enhancers, known as hypersensitive sites 1-4 (HS1-4), that drive developmental stage-specific choice and activity of the β-globin genes *HBE1*, *HBG2*, *HBG1*, *HBD* and *HBB*^47^. We verified that each of the HS1-4 produced eRNAs in divergent orientations (Fig. 5D), and hereafter refer to the HS1-4 as enhancers 1-4 (e1-4). Both eRNA and mRNA synthesis at the β-globin locus increased early in the erythroid differentiation and remained elevated upon recovery (Fig. 5D-E). Prior to their activation, the e1-4 resided in an open chromatin state and were bound by GATA1 (Fig. 5D). Remarkably, the NRF2 motif was found on each of the e1-4 enhancers (Fig. 5D), positioned upstream and proximal to the precise Transcription Start Nucleotide (TSN; Figs. 5F and S5C), which can be identified from the 5’-ends of nascent RNAs^48^. These localizations at globin enhancers agree well with the chromatin analyses (Figs. 2D and S3) and positioning of the NRF2 motifs (Fig. 5B), providing examples of NRF2 targeting the transcription initiation sites at differentiation-linked enhancers. Similarly, by analyzing the promoter region of a classical NRF2 target gene *GCLM* at nucleotide-resolution, we found the NRF2 motif directly upstream (−26 to −34 nt) of the TSN identified from the 5’-ends of nascent RNAs (Fig. S5D). These results highlight that NRF2 regulates a network of promoters and enhancers and provide a novel understanding of NRF2-driven globin transcription through the erythroid-specific enhancers.

### mRNA expression outlasts transcription

We next addressed how the differentiation-linked transcriptional changes were reflected in the steady-state mRNA expression by performing strand-specific RNA-seq after polyA-selection (mRNA-seq) (Figs. 6A and S6A-B). Due to the slow changes in the steady-state mRNA expression, we extended the early response to 12 h and the late response to 48 h. The changes in the steady-state mRNAs (Figs. 6A and S6C) were slow compared to those in nascent transcripts (Figs. 1B and 4D) but followed a similar pattern. Indeed, the transcriptionally induced NRF2 target genes were significantly induced also on mRNA level, albeit detected hours after the gene induction (Figs. 6B and S6C-D). This lag is due to the time required for the synthesis of RNA and production of mRNAs over the basal level. To report down-regulation, existing mRNAs need to be degraded, or cells proceed through mitosis to reduce the mRNA content per cell. Accordingly, the prominent downregulation of transcription (Fig. 1B) remained virtually undetected on mRNA level until 48 h (Fig. 6A), a time that comprises at least one round of cell division (Fig S2C).

**Figure 6.**
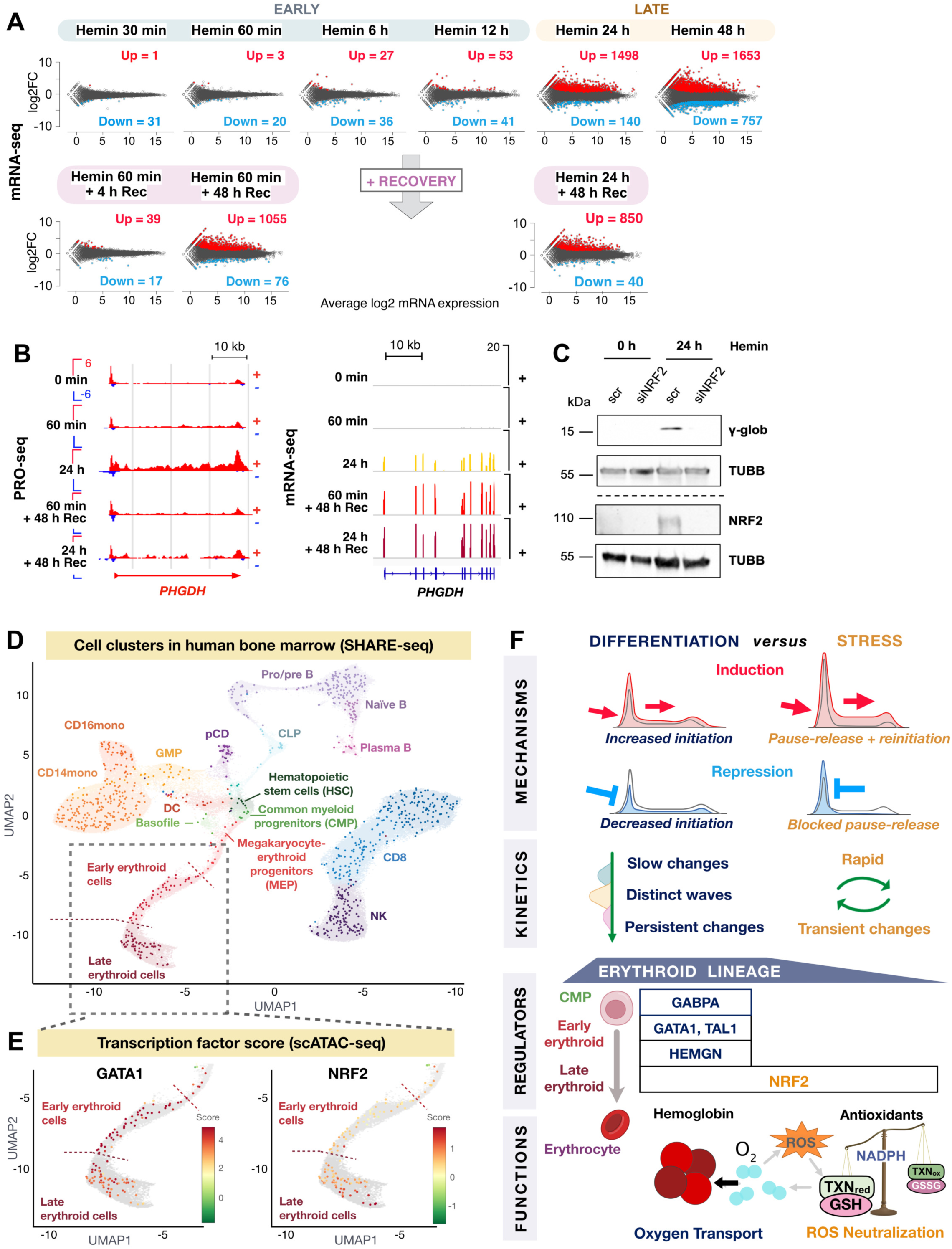
NRF2 activation occurs in late erythroid cells and in human bone marrow. **A)** Significantly (p<0.05) induced (red) or repressed (blue) mRNAs upon hemin-induced erythroid differentiation, as compared to untreated K562 cells. **B)** Nascent RNA synthesis (left) and mRNA expression (right) of phosphoglycerate dehydrogenase (*PHGDH*). **C)** NRF2 and γ-globin protein levels in K562 cells transfected with NRF2 siRNA or control siRNA (scr). β-tubulin (TUBB) serves as a loading control. **D)** Clustering of SHARE-seq (combinatorial scATAC-seq and scRNA-seq) data in human bone marrow. The distinct cell populations are indicated, and the erythroid lineage framed. **E)** Transcription factor scores of GATA1 and NRF2 during erythroid specification, showing GATA1 activity from erythroid progenitors to late erythroid cells, and NRF2 activation to peak in the late erythroid cells. The SHARE-seq data (D-E) is from reference^21^ and visualized in ACAMShiny (https://buenrostrolab.shinyapps.io/ACAMShiny/). **F)** Summary of transcription regulation in erythroid differentiation, and comparison of transcription mechanisms, kinetics, and regulators upon differentiation *versus* stress. Erythroid differentiation launches slow but persistent changes in transcription that proceed in waves and are coordinated at the rate-limiting step of initiation. On the contrary, acute stress triggers instant but transient transcriptional reprogramming, coordinated primarily at the Pol II pause-release. Differentiating cells utilize lineage-specific (blue) and stress-inducible (orange) *trans*-activators to prime and execute transcriptional reprogramming. Ordered activity of GABPA, GATA1, TAL1, HEMGN and NRF2, drive globin expression. Additionally, NRF2 activates the antioxidant program including the synthesis of glutathione (GSH), thioredoxin (TXN) and NADPH biogenesis to prepare differentiating erythroid cells to oxidative stress, encountered as the mature, enucleated oxygen-transporting erythrocytes. Related Supplementary Figures 6-8.

The overlap between transcription (PRO-seq) and mRNA expression (mRNA-seq) identified hundreds of genes that were induced or reduced on the levels of synthesis and steady-state expression (Fig. S6E). For instance, the immediate transcriptional induction of *HBG1/2* and *HBE1* (Fig. 5D-E) was followed by increased steady-state mRNA expression during the late wave (Fig. S6F). For many genes, the transcriptional activation was transient but caused long-lasting changes in the steady state mRNA expression, indicating functional consequences for the transient transcriptional induction. One of the genes that was transiently activated but caused a sustained increase in mRNA was the NRF2-regulated phosphoglycerate dehydrogenase (*PHGDH*)^49^ (Fig. 6B). PHGDH is a key enzyme in NADPH-dependent production of L-serine, a precursor in the synthesis of cysteine and glycine components of the GSH. In comparison, platelet-derived growth factor subunit B (*PDGFB*), a marker of megakaryocytic differentiation in K562 cells^50^, was downregulated throughout the early, late, and recovery-induced waves (Fig. S7A), further underscoring the hemin-induced fate decision to proceed along the erythroid lineage pathway.

### NRF2 is required for γ-globin expression

Prompted by our finding of NRF2 activity in erythroid differentiation, we examined the protein levels of NRF2 and its putative target, γ-globin (Figs. 6C and S7B). Consistent with activation of the NRF2-driven antioxidant program, NRF2 protein levels increased upon hemin-induced differentiation (Fig. S7B), showing its release from KEAP1-mediated degradation. To study whether NRF2 is required for γ-globin expression, we silenced NRF2 using RNAi and observed drastically reduced production of globin protein (Fig. 6C). These results demonstrate that NRF2 is required for an adequate production of globin, which is fundamental for the erythroid lineage, underscoring the differentiation-linked role of NRF2. Since both GATA1 and NRF2 regulate globin genes (Fig. 5), we searched the Human Protein Atlas (HPA)^51^ for their expression patterns and interaction partners. GATA1 was almost exclusively expressed in the bone marrow (Fig. S7C), whereas NRF2 was ubiquitously expressed across human tissues (Fig. S7D). The HPA reported a physical association between GATA1 and NRF2 (Fig. S7E), suggesting a direct interplay between the master regulators of differentiation and stress in erythropoiesis.

### NRF2 is activated in late erythroid cells

To couple the chromatin architecture and hemin-induced transcription kinetics to erythropoiesis *in vivo*, we used SHARE-seq^52^ data collected in human bone marrow^21^. First, we visualized hematopoietic cell clusters and differentiation paths in the bone marrow (Fig. 6D) and analyzed the accessibility of GATA1 and NRF2 motifs in the erythroid lineage (Fig. 6E). The scATAC-seq score revealed that GATA1 motifs were accessible from myeloid precursors, through the early erythroid differentiation, to the late erythroid cells (Fig. 6E, left). Instead, NRF2 motif accessibility remarkably increased at the transition from the early to the late erythroid cells and peaked at the last stages of erythroid differentiation (Fig. 6E, right). The mRNA expression of lineage-specific factors GABPA, GATA1 and HEMGN, as well as GATA1 and NRF2 target genes in human bone marrow followed the order of activation detected upon erythroid differentiation in K562 cells (Fig. S8A). These patterns of motif accessibilities and mRNA expression uncovered that hemin induced K562 cells to differentiate from multipotent myeloid progenitors to late erythroid cells. Importantly, the domains of regulatory chromatin (DORC) scores^53^, which combine the scRNA-seq and scATAC-seq in each cell, showed GATA1 activity to peak first, followed by activation of TAL1, HEMGN, and finally, NRF2 (Fig. S8B). Analyses of quantitative mass spectrometry, previously conducted in mature human erythroytes^54^, demonstrated abundant expression of globin proteins, along with proteins from the three main antioxidation pathways (Fig. S8C), underscoring the physiological importance of NRF2 activation during erythroid differentiation. Our study establishes the molecular logic of signal-induced erythroid differentiation and couples the kinetics of transcription and its regulation to *in vivo* erythropoiesis. In human, erythropoiesis is driven by propagating transcriptional waves that are coordinated by the interplay between lineage-specific transcription factors and NRF2 and produce the cellular components for oxygen transport and neutralization of ROS (Fig. 6F).

## Discussion

### Waves of gene and enhancer activity

Naïve cells acquire cell type-specific properties during differentiation by reprogramming the chromatin environment and transcription^1^. To uncover dynamics and mechanisms of transcription upon signal-induced differentiation, we tracked nucleotide-resolution changes in gene and enhancer transcription during erythroid lineage specification. RNA synthesis from genes and enhancers was reprogrammed in waves that propagated over cell divisions and altered the steady-state mRNA expression (Figs. 1, 2 and 6). In accordance with previously demonstrated hierarchical chromatin priming in hematopoiesis^55–57^, we describe an ordered production of RNA from architecturally poised enhancers and promoters. Intriguingly, a distinct set of lineage-specific transcription factors and chromatin modifiers occupied enhancers *versus* promoters (Figs. 2D and S3). Our findings underscore that the reprogramming of transcription during differentiation is primed by a promoter- and enhancer-specific chromatin environment that enables coordinated waves of RNA synthesis. Moreover, we unveil that once differentiation-linked transcriptional waves are set in motion, they cause propagating changes to the cell identity.

### NRF2 prepares for oxygen load

Cells and organisms that are exposed to mild environmental stress acquire an improved tolerance to severe challenges. This adaptive mechanism is termed hormesis and is conserved over all forms of life^58^. Stress-induced pathways can also contribute to differentiation, as shown in secreting cells that learn to tolerate a high load of protein production and stress in the endoplasmic reticulum^59^. In this study, we uncovered that erythroid differentiation prepares cells for oxidative stress that occurs in the mature erythrocytes (Figs. 4). Our data indicated that the NRF2-driven antioxidant program, which is strongly connected to alleviating elevated ROS levels, is launched in erythroid progenitors that exhibited a normal redox state. This activation of a stress response in advance is particularly interesting considering that the vital process of oxygen transport makes the enucleated erythrocytes susceptible to ROS and highly reliant on the cellular redox systems^45^. In agreement, erythrocytes in Nrf2-deficient mice are morphologically abnormal, decreased in number, and susceptible to oxidative damage^60^. We propose that NRF2 plays an integral part of erythroid differentiation by interacting with lineage-specific transcription factors (Figs. 5 and S7) and building up reservoirs of GSH, TXN and enzymes for NADPH regeneration (Fig. 4), preparing erythroid progenitors for their mature role as oxygen carriers (Fig. 6F).

### Molecular control of differentiation

Since erythroid differentiation activated a stress-induced transcription factor, we directly compared mechanisms that reprogram transcription upon differentiation and stress. The kinetics and the steps of transcription that coordinated Pol II molecules were drastically different in erythroid differentiation *versus* acute stress (Figs. 3 and 6F). Transcriptional responses to both heat shock^15^ and oxidative stress^18^ are primed by paused Pol II, and the paused Pol II instantly launched into productive elongation at induced genes. Instead, the stress-repressed genes stall Pol II at the pause (Fig. 3)^15,18^, which effectively bookmarks genes for rapid reactivation without the need to close or remodel the chromatin^16^. In stark contrast, transcriptional reprogramming upon erythroid differentiation was coordinated primarily at the initiation and caused long-lasting changes in the transcription program (Fig. 3). Our results are well-aligned with studies that identified transcription initiation as the principal regulatory step in mouse erythropoiesis^61^, and a rapid transcriptional response in macrophages by releasing paused Pol II^42^. Together, these genome-wide analyses of transcriptional reprogramming demonstrate Pol II pausing to enable rapid, synchronous, and transient responses, while redistributing transcription machinery across genes and enhancers *via* initiation causes slow but long-term transcriptional changes. These mechanisms of genome-wide coordination of Pol II molecules uncover strategies by which cells prime and execute transcriptional reprogramming upon distinct conditions (Fig. 6F).

### Interplay of master regulators

It has been previously established that NRF2 regulates γ-globin mRNA expression by binding to the γ-globin promoter and enhancers within the LCR^62^. Our study expanded the role of NRF2 in erythroid differentiation by demonstrating the induction of a comprehensive network of NRF2 target genes (Fig. 4) and enhancers (Fig. 5). By closely examining the erythroid-specific enhancers in the β-globin LCR, we found that NRF2 motifs co-localized with GATA1-binding sites, suggesting that NRF2 drives globin genes in collaboration with the erythroid-specific master regulator. NRF2 has been reported to interact with GATA1 *via* the Neh6 domain^63^ that supports NRF2 degradation even in the absence of KEAP1^64^. It is tempting to speculate that upon erythroid differentiation, GATA1 could stabilize NRF2 by inhibiting the access to the Neh6 domain, activating NRF2 in the absence of elevated levels of ROS. Moreover, GATA1 could bookmark promoters and enhancers for NRF2-driven activation, and recruit NRF2 *via* direct binding. We propose that lineage-specific transcription factors prime promoters and enhancers for prompt activation upon differentiation, and the potent stress regulators, including NRF2, are utilized as rapid *trans*-activators to elicit signal-induced transcriptional changes. In erythropoiesis, the interplay between master regulators of differentiation and stress drives lineage specification and produces the cell type-specific machineries for oxygen transport and ROS neutralization (Fig. 6F).

## Resource availability

### Lead contact

Further information and requests for resources and reagents should be directed and will be fulfilled by the Lead Contact, Assist. Prof. Anniina Vihervaara.

### Materials availability

This study did not generate new unique reagents.

### Data and code availability

The PRO-seq and mRNA-seq datasets generated in this study have been deposited to GEO (https://www.ncbi.nlm.nih.gov/geo/). Original figures of Western Blotting presented in this study are available in Mendeley. Computational analyses have been conducted using shell/bash, R and Python, and the main pipelines for mapping PRO-seq and mRNA-seq data (https://github.com/Vihervaara/PRO-seq-analyses; https://github.com/Vihervaara/mRNA-seq) and identification of functional genomic regions (https://github.com/Vihervaara/functionalGenomicRegions) are available in GitHub. Custom made script for downstream analyses can be made available from the lead contact.

## Material and methods

### Cell culture and hemin treatment

Human myeloid leukemia K562 cells originated from ATCC, were tested to be mycoplasma free, and verified by imaging, cell cycle analyses and γ-globin expression to display the morphology and characteristics of K562 cells. The cells were maintained at 37°C in a humidified 5% CO_2_ atmosphere and cultured in RPMI 1640 medium (Sigma-Aldrich) supplemented with 10% (vol/vol) fetal bovine serum (FBS), 2 mM L-glutamate, 100 μg/ml streptomycin, and 100 U/ml penicillin. Hemin was dissolved in 1 M NaOH to obtain a 10 mM hemin stock solution. In all experiments, cells were treated with 30 μM hemin. After the hemin treatment, cells were either directly collected, or thoroughly washed and allowed to recover in hemin-free media.

### Cell viability and cell proliferation

Cells were counted and their viability assessed during hemin treatments and recovery with automated cell analyzer NucleoCounter NC-250 (ChemoMetec). The relative change in cell count is indicated as [number of cells after recovery/number of cells after treatment]. The viability was above 95% at all conditions, and the relative change in cell viability is indicated as [viability after recovery/viability after treatment]. The error bars indicate standard deviation.

### Cell cycle analysis

DNA content in hemin-treated K562 cells was determined by flow cytometry with propidium iodide (PI) staining. For the analysis, 0.5 million cells were fixed in ice-cold 70%-EtOH. The EtOH-fixed cells were collected by centrifugation at 700 x g for 5 min and washed once in PBS + 1%-FBS. The cell pellet was resuspended in FxCycle PI/RNase Staining Solution (Invitrogen, Thermo Fisher Scientific, Inc) and the dye was incubated for 30 min at room temperature (RT). The fluorescence-mediated cell counting was performed using the LSRFortessa Cell Analyzer (BD Biosciences) flow cytometer with the BD FACSDiVa software v.8. The FACS histograms were generated using FlowJo v.10 (BD Biosciences).

### Transient transfections

Transient transfections of K562 cells were performed using the Neon Transfection System (Invitrogen, Thermo Fisher Scientific) with the conditions of 1,450 V, 10 ms, and 3 pulses. To downregulate NRF2 expression, ON-TARGETplus Human NFE2L2 siRNA SMARTPool and ON-TARGETplus Non-targeting Control Pool (Horizon Discovery, Perkin Elmer) were used. Transfected cells were allowed to recover for 24 h before hemin treatments.

### Chromatin isolation for PRO-seq

PRO-seq was performed in untreated (0 min) cells, in cells treated with 30 μM hemin for 15 min, 30 min, 60 min or 24 h, and in cells that had recovered from 60 min or 24 h hemin for 48 h. Instantly after each treatment, cells were placed in excess of ice-cold PBS, washed twice with the PBS and the chromatin isolated in NUN buffer (0.3 M NaCl, 1 M urea, 1% NP-40, 20 mM HEPES pH 7.5, 7.5 mM MgCl_2_, 0.2 mM EDTA, 1 mM DTT), complemented with 20 units SUPERaseIn RNase Inhibitor (Life Technologies # AM2694) and 1x cOmplete EDTA-free Protease Inhibitor Cocktail (Roche cat nr. 11873580001). The samples were thoroughly vortexed and centrifuged at 12,500 x g at 4°C for 30 min. The chromatin pellets were washed in ice-cold 50 mM Tris-HCl (pH 7.5) and stored at −80°C in chromatin storage buffer (50 mM Tris-HCl pH 8.0, 25% glycerol, 5 mM MgAc_2_, 0.1 mM EDTA, 5 mM DTT, 20 units SUPERaseIn RNase Inhibitor). The chromatin was sonicated for 5 min using Bioruptor (Diagenode) with 30 s on / 30 s off intervals and high throughput settings.

### Run-on reaction and nascent RNA isolation

Chromatin from 10 million K562 cells was used per run-on reaction, and the run-on conducted in the presence of 25 μM biotin-11-A/C/G/UTP (Perkin Elmer cat nrs. NEL544001EA, NEL542001EA, NEL545001EA, NEL543001EA), in 5 mM Tris-HCl pH 8.0, 2.5 mM MgCl_2_, 0.5 mM DTT, 150 mM KCl, 40 units SUPERaseIn RNase inhibitor and 1% sarcosyl. Total RNA was isolated using TRIzol LS and chloroform, and EtOH precipitation containing glycoblue (ThermoFisher, AM9516) as a marker of the precipitate. The RNA was base hydrolyzed with NaOH for 5 min, and the base hydrolysis terminated with Tris-HCl pH 6.8. The samples were passed through P-30 columns (BioRad, cat.nr. 7326232) to remove free biotinylated nucleotides, and the nascent RNAs, containing biotin-nt at the 3’-end, purified from the non-nascent RNAs using Streptavidin Dynabeads (Invitrogen, MyOne C1, cat. nr. 65002). The bead-bound nascent-RNAs were washed with wash buffers 1 (50 mM Tris-HCl pH 7.4, 2 M NaCl, 0.5% Triton X-100), 2 (10 mM Tris-HCl pH 7.4, 300 mM NaCl, 0.1% Triton X-100) and 3 (5 mM Tris-HCl pH 7.4).

### PRO-seq library preparation

The nascent RNA was isolated from the beads with TRIzol and chloroform, and precipitated with EtOH using glycoblue as the marker. The RNA pellet was re-diluted and barcoded 3’-adapters ligated over night at 25°C using T4 RNA Ligase I (NEB, M0204L) in 10 µl reaction volume and 5 µM final 3’-adapter concentration. The sample-specific barcoded 3’-adapters contained a constant G at the ligation site (5’-end of the adapter), followed by a 6-nt in-line barcode, and an inverted T at the 3’-end to prevent adapter concatemers. The unligated adapters were removed by binding the nascent transcripts to Streptavidin Dynabeads and washing the beads with wash buffers 1 and 3. During the last wash, the barcoded samples were combined (a maximum of four samples pooled per tube) for eased handling. Next, the 5’-end of nascent RNAs was decapped on-beads in RNA 5’ Pyrophosphorylase mix (NEB, M0356S) at 37 °C for 45 min, followed by 5’-hydroxyrepair in T4 polynucleotide kinase mix (NEB, M0201S) at 37 °C for additional 45 min. The bead-bound nascent RNAs were washed with buffers 1 and 3, and the nascent transcripts isolated from the beads with TRIzol and chloroform, and precipitated with EtOH. The air-dried RNAs were ligated to UMI-containing 5’-adapters with T4 RNA Ligase I over night at 25°C in 20 µl reaction volume and 2.5 µM final 5’-adapter concentration. The 5’-adapter contained a constant C at the ligation site (3’-end of the adapter), an adjacent 6-nt UMI, and an inverted T at the 5’-end to prevent adapter concatemers. Unligated adapters were removed by binding the nascent transcripts to Streptavidin Dynabeads and washing the bead-nascent-RNA-complexes with wash buffers 1, 2 and 3.

### Reverse transcription and final amplification

The nascent transcripts were isolated with TRIzol and chloroform and precipitated with EtOH. Reverse transcription was performed with Super Script III reverse transcriptase (ThermoFisher, 18080044) in 25 µl reaction volume containing 2.5 µM RP1-primer, 625 µM dNTPs and 20 units SUPERaseIn RNA inhibitors using the following settings: 15 min at 45°C, 40 min at 50°C, 10 min at 55°C, 15 min at 70°C. After test-amplification to verify the library qualities, the samples were amplified in a total of 11 cycles using Phusion Polymerase (NEB, M0530S) in 1x HF buffer, 1 M betaine, 0.25 mM dNTPs, 0.2 µM RP1 primer and 0.2 µM RPI-n index primer. The amplified samples were purified with 1.6x Ampure XP beads (Beckman Coulter, cat. nr. A63880), analyzed with BioAnalyzer and sequenced with Illumina NovoSeq6000 (NovoGene Inc, CA, USA) using PE150 settings. RP1 and RPI-n primers are Illumina small RNA TruSeq design (Oligonucleotide sequences © 2015 Illumina, Inc. All rights reserved).

### Data analyses of PRO-seq libraries

The computational analyses follow the established pipelines for mapping (https://github.com/Vihervaara/PRO-seq-analyses)^16,48^ and functional regulatory element identification (https://github.com/Vihervaara/functionalGenomicRegions)^32^. In brief, reads in a pool of samples were first separated using fastx barcode splitter (http://hannonlab.cshl.edu/fastx_toolkit), after which, correct pairing of reads was ensured with fastq_pair (https://github.com/linsalrob/fastq-pair). The base adapter sequences were removed, and the 6-nt UMI in the 5’-adapter was identified using fastp^65^. The 7-nt barcode+C and 7-nt UMI + C were then removed and the read 1 reverse complemented with fastx toolkit. The read pairs were mapped to the human genome (hg19) with Bowtie2, and PCR duplicates collapsed using UMI tools^66^. The collapsed bam files were processed to bed, bedgraph and bigwig files using samtools^67^, bedtools^68^, and bedGraphToBigWig (https://www.encodeproject.org/software/bedgraphtobigwig/). The complete PRO-seq datasets with raw (fastq) and processed (bigWig) files are available via Gene Expression Omnibus database (https://www.ncbi.nlm.nih.gov/geo/).

### Quantification of engaged Pol II at genes and enhancers

In PRO-seq, a single biotinylated nucleotide is incorporated into the 3’-end of a growing RNA chain, enabling isolation of nascent transcripts from the total pool of RNA, and mapping the active sites of transcription at nucleotide-resolution genome-wide^32,35^. In mammals, promoters and enhancers contain two core initiation regions that recruit the preinitiation complex (PIC) and position Pol II to the transcription start site^46^. This promoter and enhancer architecture generates a characteristic pattern of divergent transcription that can be utilized to identify the actively transcribed genes and enhancers in the whole genome^15,32,69^. We used the pattern of divergent transcription to identify active genes and enhancers^32,69^ in each of the PRO-seq dataset generated. The coordinates of active enhancers across the individual samples were merged using bedtools merge -d -100 to generate unified enhancer coordinates across the samples. To quantify enhancer transcription, Pol II density was quantified strand-specifically along the enhancer. Gene activity was measured from the gene body (+500 from the annotated TSS to −500 from the annotated cleavage and polyadenylation site, CPS) as previously described^15,32^. Pol II density at promoter-proximal region was quantified from 50 nt window with the highest Pol II density, searched from −100 to +400 from the annotated TSS. Hemin-induced changes in gene or enhancer transcription were analyzed with DESeq2^70^ against untreated (hemin 0 min) sample using adjusted p-value 0.01 and minimum absolute fold change 1.25.

### Strand-specific mRNA-seq

K562 cells were grown to a concentration of 0.5 million cells/ml and mRNA-seq samples prepared in three independent replicates. Cells treated with 30 μM hemin were harvested at 0, 30 and 60 min and 6, 12, 24 and 48 h by washing twice with ice-cold PBS (BioWest) and flash-freezing. Recovery samples were prepared after 60 min or 24 h of hemin treatment (60 min + 4 h rec; 60 min + 48 h rec; 24 h +48 h rec) as described above. The total RNA was isolated with, AllPrep DNA/RNA/miRNA Universal Kit (Qiagen), alternatively by purification with Trizol and chloroform, followed by EtOH precipitation.

From the total RNA, polyA-containing RNA (mRNA) was isolated and a cDNA library constructed using Stranded mRNA Prep kit as described by the manufacturer (Illumina). The quality of the cDNA library was controlled at the Finnish Functional Genomics Centre (Turku Bioscience) by using the 5300 Fragment Analyzer system (Agilent Technologies Inc). The cDNA libraries were sequenced at the Novogene US Davis Sequencing Center (Sacramento, CA, USA) using the Illumina NovaSeq 6000 platform with a PE150 strategy. The raw fastq files and normalized density profiles are available in GEO.

### Computational analyses of mRNA-seq data

The obtained fastq files were trimmed with fastx_tool and mapped to hg19 with Hisat2^71^. Reads at each gene were counted with featureCounts^72^ opting for strand-specificity and using gencode.v19.annotation.gtf (https://www.gencodegenes.org/human/release_19.html) file as the reference. The correlation of the replicates was verified and differential expression analysis between hemin and control condition were performed using DEseq2 using p-adjusted 0.05, minimum absolute fold change 1.25 and minimum of mean reads across replicates per gene 10.

### Visualization of engaged Pol II, mRNA expression and chromatin state

The PRO-seq, mRNA-seq, ChIP-seq and DNase I-seq datasets were visualized using an in-house tool (Hojoong Kwak, Cornell University, Ithaca, USA) and Integrative Genomics Viewer (IGV)^73^. The scale of the y-axis is indicated on the top browser track for each panel and is the same and linear for all tracks in the browser image. The signal on the plus strand is shown on a positive scale, the signal on the minus strand is shown on a negative scale. The metaprofiles show the average density for the indicated data at genes or enhancers. The shaded areas indicate a confidence interval between 12.5% - 87.5%. NRF2 motifs in genome browser are: TGACnnnGC sequences; GATA motifs: GATA sequences; and the initiator (Inr): CA dinucleotide when the A overlaps a precise +1nt identified from the 5’-ends of nascent RNAs^48^ and the C is at the −1 position.

### Gene Ontology, Enrichr and MEME-ChIP analyses

Gene sets obtained from PRO-seq and mRNA-seq analysis were examined for enriched functionalities, transcription factor motifs and transcription factor regulation. GO analyses were conducted with GO resource with the PANTHER 18.0 annotation version^74–76^ (date of analysis October 2, 2023). To make predictions of transcriptional regulators of the obtained gene sets, we used the Enrichr bioinformatics tool^77^ with the ChEA Consensus TFs from ChIP-X database (date of analysis October 3, 2023). To analyze enrichment for transcription factor binding sites we utilized MEME-ChIP^78^ with HOCOMOCOv11_full and settings for -minw 5 - maxw 20 -nmotifs 10. As input data we analyzed a 1,000 nt-region centered to the promoter TSS or midpoint of enhancers. The probability of obtained motifs with respect to the enhancer center was analyzed by CentriMo 5.5.5^79^. In all above bioinformatics analyses we regarded findings with p<0.05. The occurrence of NRF2 motif on hemin-induced enhancers were compared to the respective occurrence at unchanged enhancers. The following steps were performed for all time points: first, genomic coordinates of the optimal^80^ NRF2 motif “TGACTCAGC” occurrences in hg19 were generated as a bed file, utilizing fastaRegexFinder (https://github.com/dariober/bioinformatics-cafe/blob/master/fastaRegexFinder.py).

Second, coordinates of regulatory regions were intersected with the coordinates of the optimal NRF2 motifs using bedtools intersect^68^ (version 2.31.0). The fraction of regulatory regions containing at least one NRF2 motif was calculated for hemin-induced and unchanged regions separately. Subsequently, a chi-square test for independence was performed on a contingency table with the four following categories: hemin-induced/unchanged regulatory regions, and regulatory regions with/without at least one NRF2 motif.

### Tissue-specific mRNA expression and protein interaction analyses

The HPA^51^ was used to investigate mRNA expression patterns across human tissues and protein interactions between GATA1 and NRF2. The HPA links are as follows: https://www.proteinatlas.org/ ENSG00000102145-GATA1/interaction, https://www.proteinatlas.org/ENSG00000116044-NFE2L2/interaction, https://www.proteinatlas.org/ENSG00000102145-GATA1/tissue; https://www.proteinatlas.org/ENSG00000116044-NFE2L2/tissue.

### GSH/GSSG assay

The amounts of reduced (GSH) and oxidized (GSSG) glutathione in K562 cells upon erythroid differentiation and oxidative stress were measured using GSH/GSSG-Glo Assay (Promega), which utilizes luciferase for glutathione detection in cell lysates. For erythroid differentiation, cells were treated with 30 µM hemin for 60 min and 24 h. The two different treatments were started so that they ended concurrently. For oxidative stress, K562 cells were treated with 10, 50, 75, and 100 µM menadione (MD) for 60 min. Luminescence was measured using the Hidex Sense microplate reader (Hidex). The experiments were performed in triplicate. The amount of GSH was determined by subtracting GSSG from the total amount of glutathione, and GSSG by dividing the relative luminescence of GSSG by 2, according to the manufacturer’s instructions. The relative amounts of GSH and GSSG were determined by normalizing to the corresponding form of glutathione in the basal condition. To determine the difference between two groups, unpaired Welch’s unequal variances t-test was applied. Error bars in the statistical analyses indicate standard deviations.

### Immunoblotting

Cells were collected, washed with PBS (BioWest) and lysed in Laemmli buffer with 3% β-mercaptoethanol. Samples were denatured at 95°C for 5 min, loaded onto a gradient 4%-20% or 7.5% Mini-PROTEAN TGX precast gel (Bio-Rad), and transferred to nitrocellulose membranes using the Trans-Blot Turbo Transfer System (Bio-Rad). For NRF2 detection, membranes were blocked in 5% milk-TBS-0.01% Tween20 for 60 min at RT. The NRF2 antibody (D1Z9C, Cell Signaling Technology) was diluted in 5% milk-TBS-0.1% Tween20, applied to the membranes, and the membranes were incubated overnight at 4°C. For the other antibodies, an HRP-conjugated anti-globin γ (51-7 HRP, sc-21756, Santa Cruz Biotechnology) and anti-β-tubulin (T8328, Sigma-Aldrich), membranes were blocked in 5% milk-PBS-0.03% Tween20 solution, and the primary antibodies diluted in 0.5% BSA-PBS-0.02% NaN3 and incubated on membranes overnight at 4°C. HRP-conjugated rabbit and mouse secondary antibodies (Promega) were used to detect anti-NRF2 and anti-β-tubulin, respectively. Amersham ECL Western Blotting Detection Reagent (Cytiva) or the SuperSignal West Pico or Femto Chemiluminescence Substrate kits (Life Technologies) were used for detection. All immunoblotting experiments were performed at least three times. The original uncropped WB images have been deposited to Mendeley (private state until publication, link provided upon request).

### SHARE-seq data analyses

Count matrices and metadata of scRNA-seq and scATAC-seq, originating from SHARE-seq in human bone marrow cells^21^, was downloaded from the GEO (GSE216461). The metadata was utilized to match scRNA-seq and scATAC-seq by each cell, and the cell types (clusters) sorted from the column “cistopic.assign.l2”. The cell types in the lineage from the hematopietic cell to late erythoid cells (HSC/MPP, CMP, MEP, early-Ery, late-Ery) were selected and the distribution of mRNA expression analysed in each group. To visualise the clusters of human bone marrow cells, including ATAC-seq scores and DORC scores in the erythroid lineage, the shinyapp from Buenrostro lab was used (https://buenrostrolab.shinyapps.io/ACAMShiny/). Transcription factor scores for GATA1 and NRF2 are shown in the figure. The DORC scores are as adjusted by the ACAMShiny when all bone marrow cells are shown (date of analyses, June 2^nd^, 2024).

### Quantification of protein expression in mature human erythocytes

Peptide counts in mature human erythrocytes were obtained as a matrix^54^. The total sum of peptides was plotted for all identified proteins, NRF2 target genes from the three antioxidation pathways, and globin proteins.

### Schematic figures

Schematic figures were created with Keynote and AffinityDesigner.

## Additional datasets used

Besides the PRO-seq and mRNA-seq datasets generated in this study, the following datasets have been utilized: ChIP-seq and DNase I-seq in non-stressed K562 cells were obtained from the ENCODE^36^ (http://encodeproject.org/); PRO-seq data in non-stressed and heat-shocked K562 cells were downloaded from GEO using accessions GSE89230 [ref 15], GSE127844, GSE154746 and GSE128160 [ref 16]. SHARE-seq data, containing scRNA-seq and scATAC-seq in human bone marrow, was obtained from GEO using accession GSE216464 and its subseries GSE216461 and GSE216461, originating from [ref 21]. Quantitative mass spectrometry in human erythrocytes was obtained from [ref 54].

## Acknowledgements

We thank the members of the Vartiainen, Sistonen and Vihervaara laboratories for valuable discussions during the manuscript preparation. Ville Koponen is acknowledged for expert assistance with protein analyses. This work was financially supported by Swedish Research Council (A.V. grant number 2021-02668), SciLifeLab and KTH (A.V. SciLifeLab Fellowship), Research Council of Finland (A.V. grant number 32495; L.S. grant number 355596; M.V. grant numbers 330254 and 338281), Sigrid Jusélius Foundation (L.S. and M.V.), Cancer Foundation Finland (L.S.) and Åbo Akademi University (L.S.).

## Author contributions

L.S. and A.V. conceived and designed the study. I.K., J.C.P., S.V.H, B.P., E.B.J., and A.V. conducted the laboratory work, and I.K., A.R., S.V.H., and A.V. performed the computational data analyses. All the authors interpreted the results and contributed to the study outline. M.K.V., L.S. and A.V. supervised the work. I.K., L.S., and A.V. wrote the manuscript with input from J.C.P., A.R., S.V.H., B.P. E.B.J., and M.K.V.

## Supplementary material

Datasets:

**Dataset 1:** List of differentially expressed genes and enhancers upon hemin treatment of K562 cells measured with PRO-seq and mRNA-seq analyses.

**Dataset 2:** Gene ontology (GO) term enrichment in hemin-induced genes (Gene Ontology Resource)

**Dataset 3:** Transcription factor target gene enrichment among differentially expressed genes detected with PRO-seq (Enrichr)

**Dataset 4:** Motif occurences on hemin-induced enhancers (MEME-ChIP)

## Supplementary figure legends

**Supplementary Figure 1.**
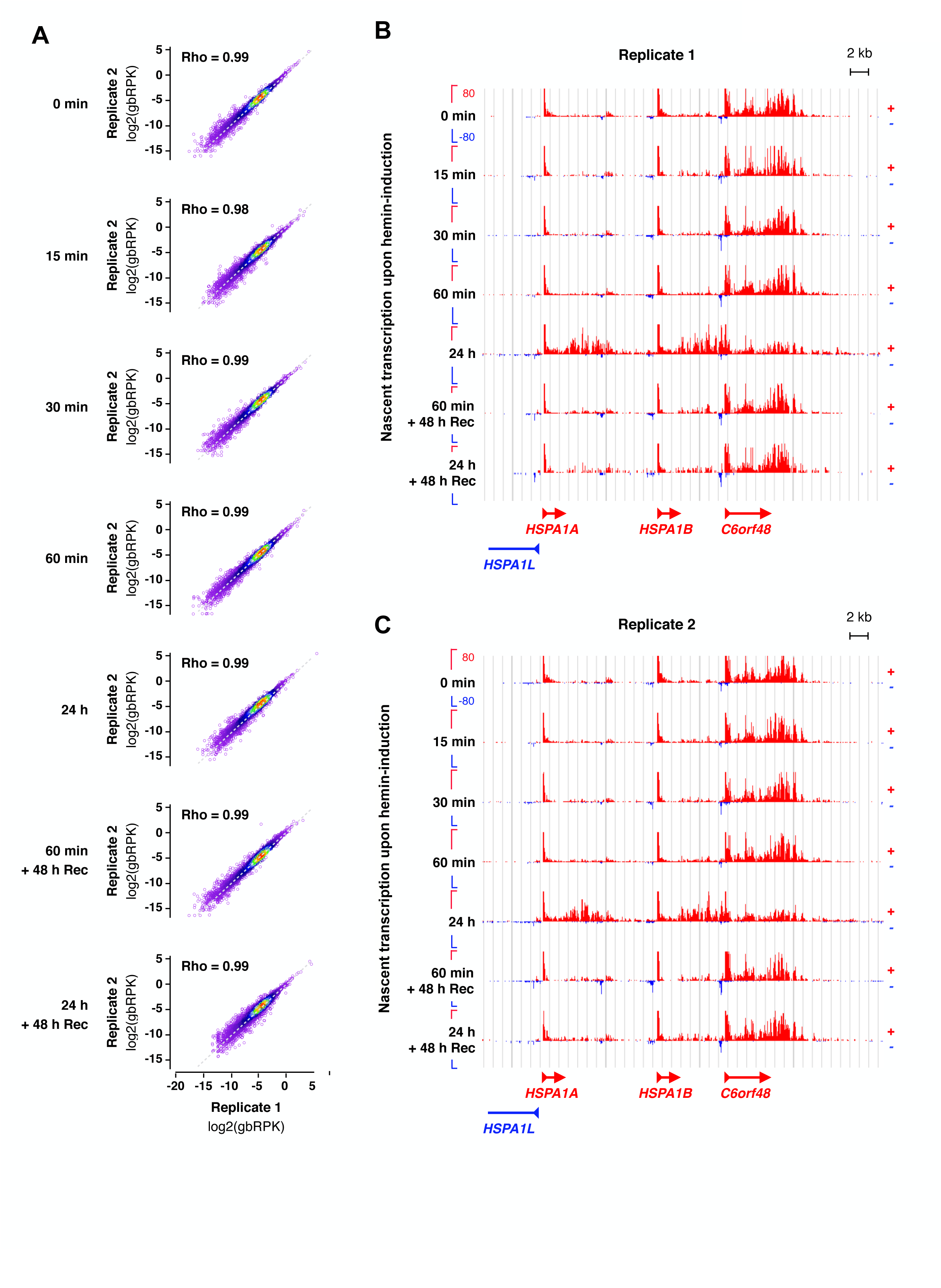
PRO-seq replicates show high correlation. **A)** Scatterplots of nascent transcription at the gene body (gb) of each gene in replicate 1 (x-axis) and replicate 2 (y-axis). Rho indicates Spearman’s rank correlation. The dotted gray line indicates x=y. **B-C)** Genome browser images of transcription at the *Hsp70* locus in replicate 1 (B) and replicate 2 (C)

**Supplementary Figure 2.**
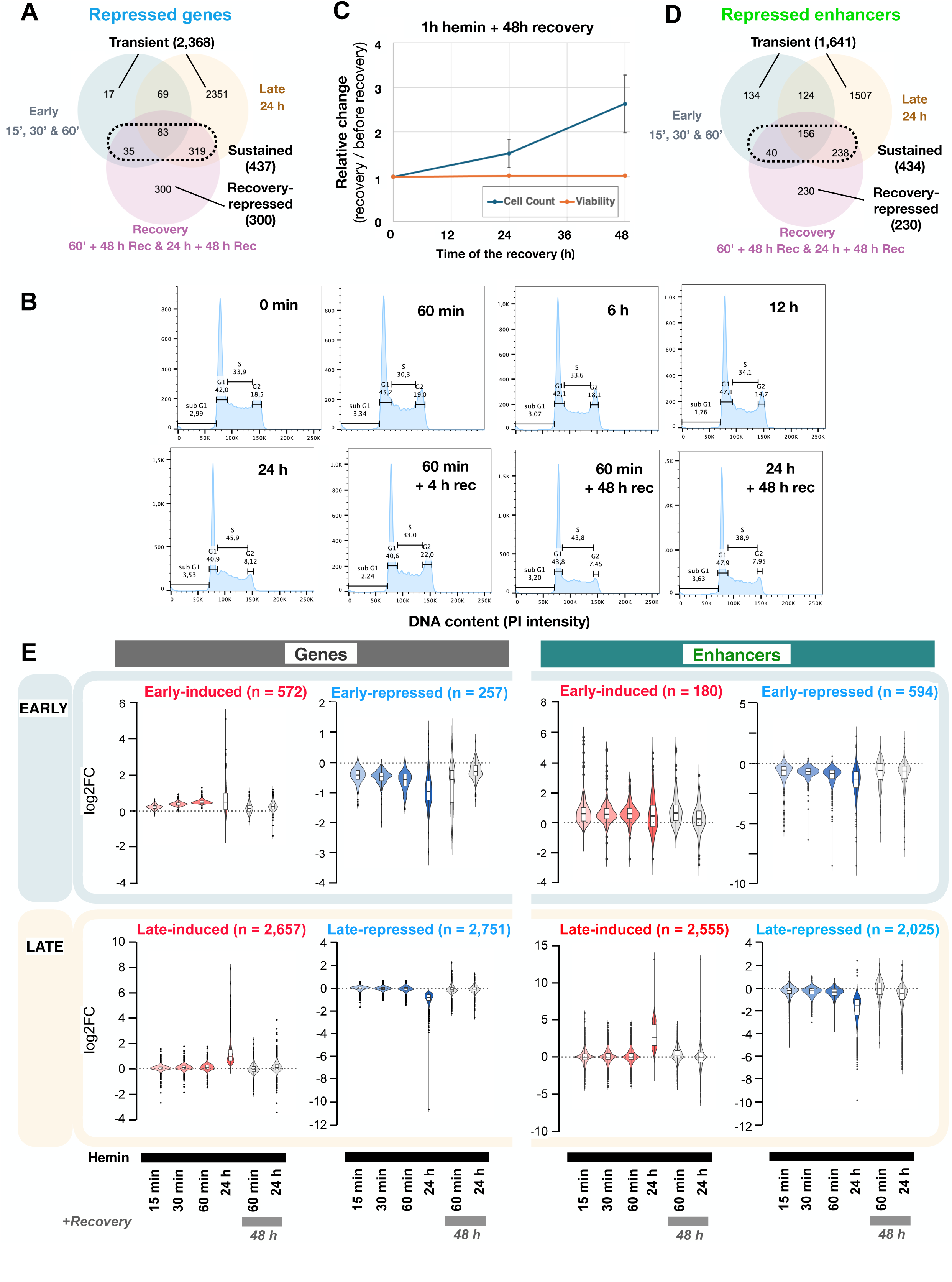
Early-induced changes in gene and enhancer transcription are carried over cell divisions. **A)** Overlap between early-, late-, and recovery-repressed genes. **B)** Cell cycle profiles of hemin-treated K562 cells, analyzed by propidium iodide (PI) staining of the DNA content using flow cytometry. **C)** Cell proliferation and viability measured during recovery from a 60-min hemin induction. The relative cell count is given as [cell number after 48-h recovery / cell number before recovery]. The relative cell viability is given as [fraction viable cells after 48-h recovery / fraction viable cells before recovery]. The analyses of cell proliferation and viability were conducted in three replicates and the error bars indicate standard deviation. **D)** Overlap between early-, late-, and recovery-repressed enhancers. **E)** Violin profiles of transcriptional changes at induced and repressed genes (leftmost panels) and enhancers (rightmost panels).

**Supplementary Figure 3.**
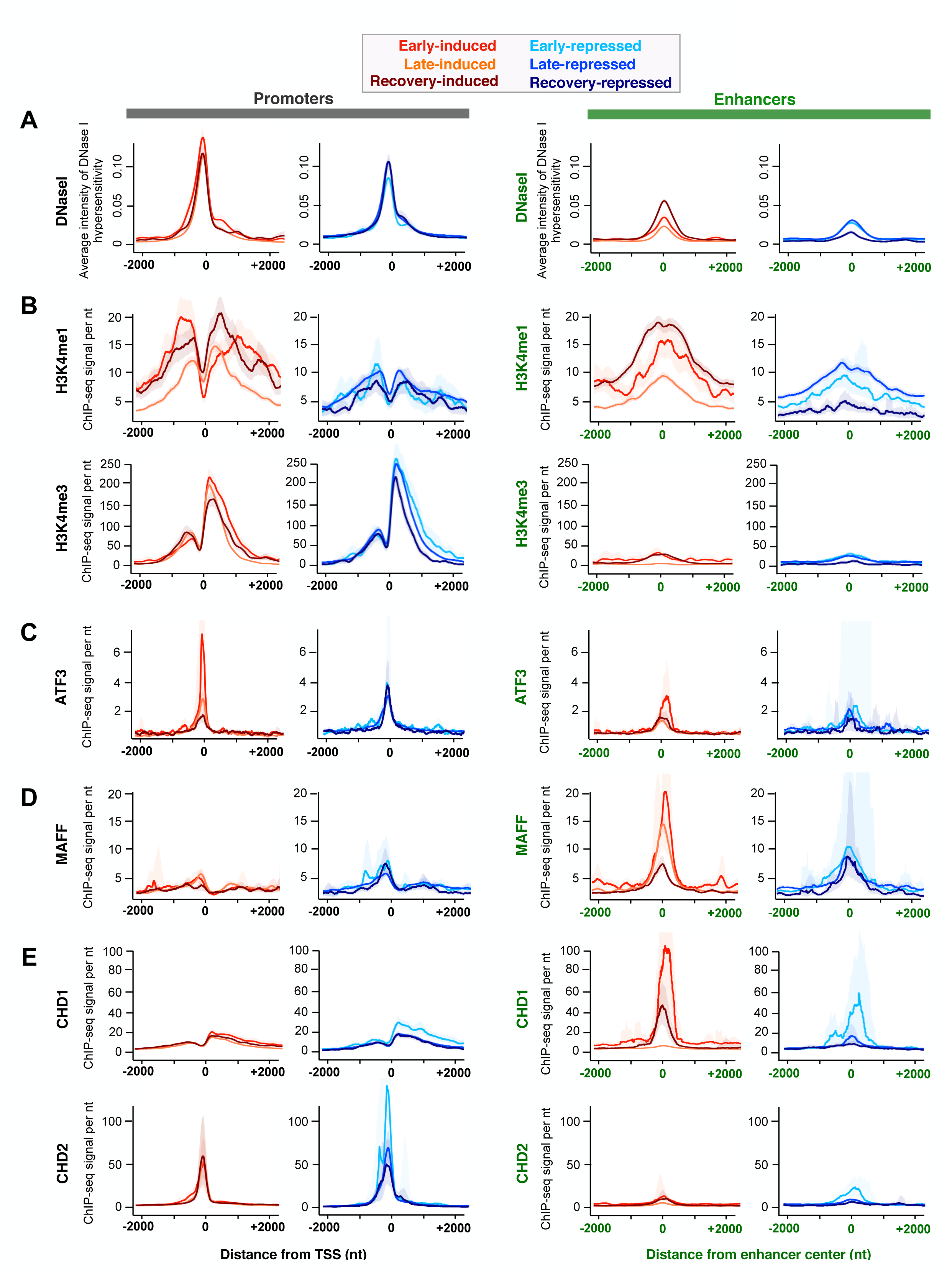
Chromatin architecture primes promoter and enhancer activation during erythroid differentiation. **A)** Average intensity of DNase I hypersensitivity along hemin-induced and repressed regulatory elements. **B-E)** Average ChIP-seq intensity of **B)** histone modifications H3K4me1 and H3K4me3, **C)** transcription factor ATF3 **D)** small Maf protein MAFF, and **E)** chromatin remodelers CHD1 and CHD2, at hemin-induced and repressed promoters and enhancers. The shaded area around the average denotes 12.5-87.5% confidence interval. The DNAse I-seq and ChIP-seq data are from the ENCODE^36^.

**Supplementary Figure 4.**
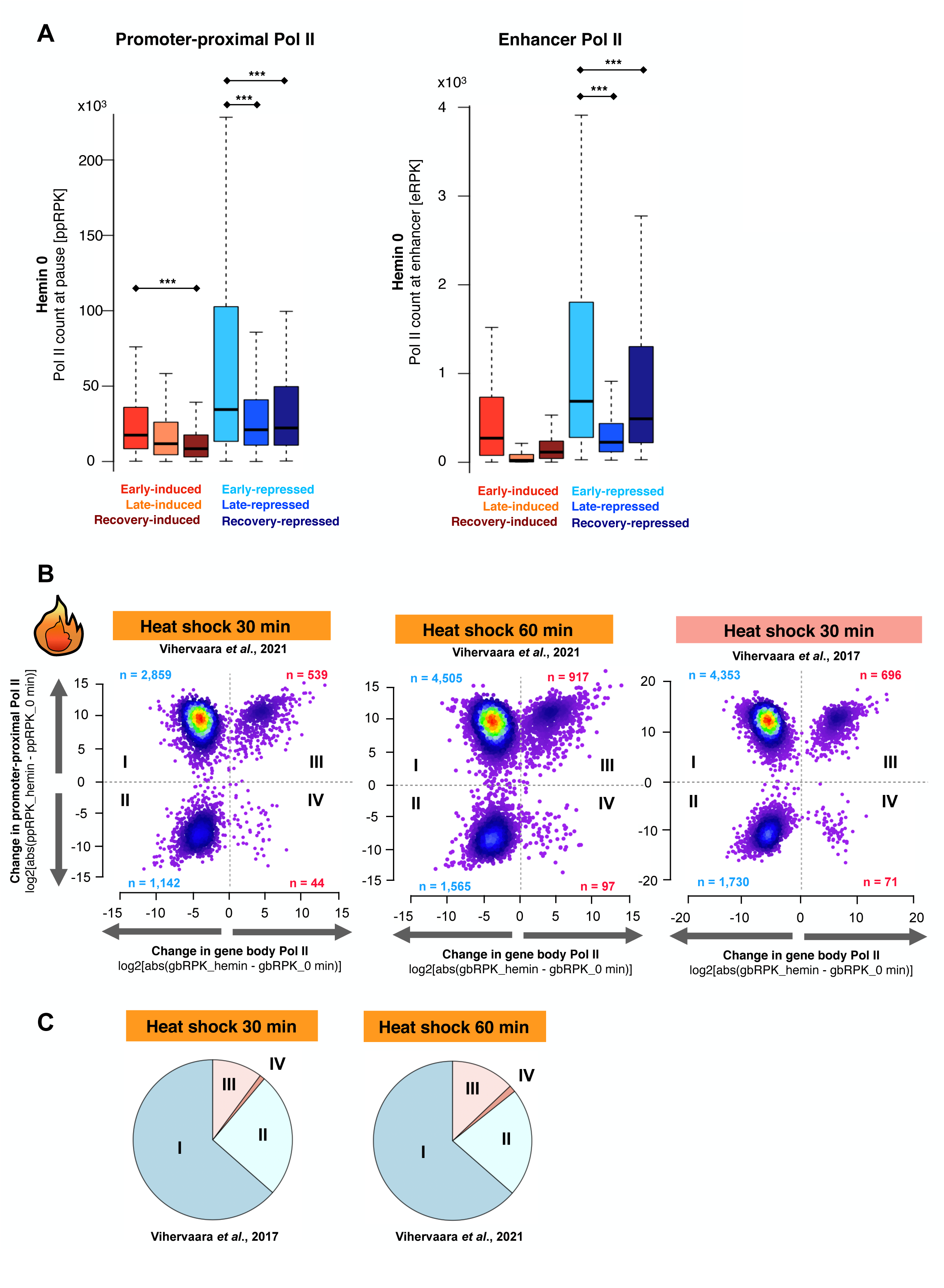
Heat shock induces instant transcriptional repression by inhibiting Pol II pause-release. **A)** Quantification of paused Pol II at promoter-proximal region of hemin-induced and repressed genes (left panel), and engaged Pol II at hemin-induced and repressed enhancers (right panel), before the hemin-treatment. The statistical analyses were conducted with ANOVA. ***p-value < 0.001. **B)** Scatter plots deciphering the change in Pol II occupancy at the promoter-proximal pause region and the gene body upon heat shock. Each gene is indicated with a sphere and positioned based on the change [heat shock - control] in Pol II pausing (y-axis) and productive elongation (x-axis). Genes in the top left square (I) are repressed by inhibiting the release of Pol II into productive elongation (increased Pol II pausing and reduced elongation). In the bottom left square (II) genes are repressed by decreased transcription initiation (less Pol II at the pause region and along the gene body). Genes in the top right square (III) are induced by an increased rate of transcription initiation (increased Pol II at the pause region and along the gene). In the bottom right square (IV), the rate of pause-release increases more than the rate of initiation (reduced pausing, increased elongation). The PRO-seq data originates from references^15–16^. **C)** Proportion of heat-induced and heat-repressed genes in the categories I-IV upon 30-minute and 60-min heat treatments.

**Supplementary Figure 5.**
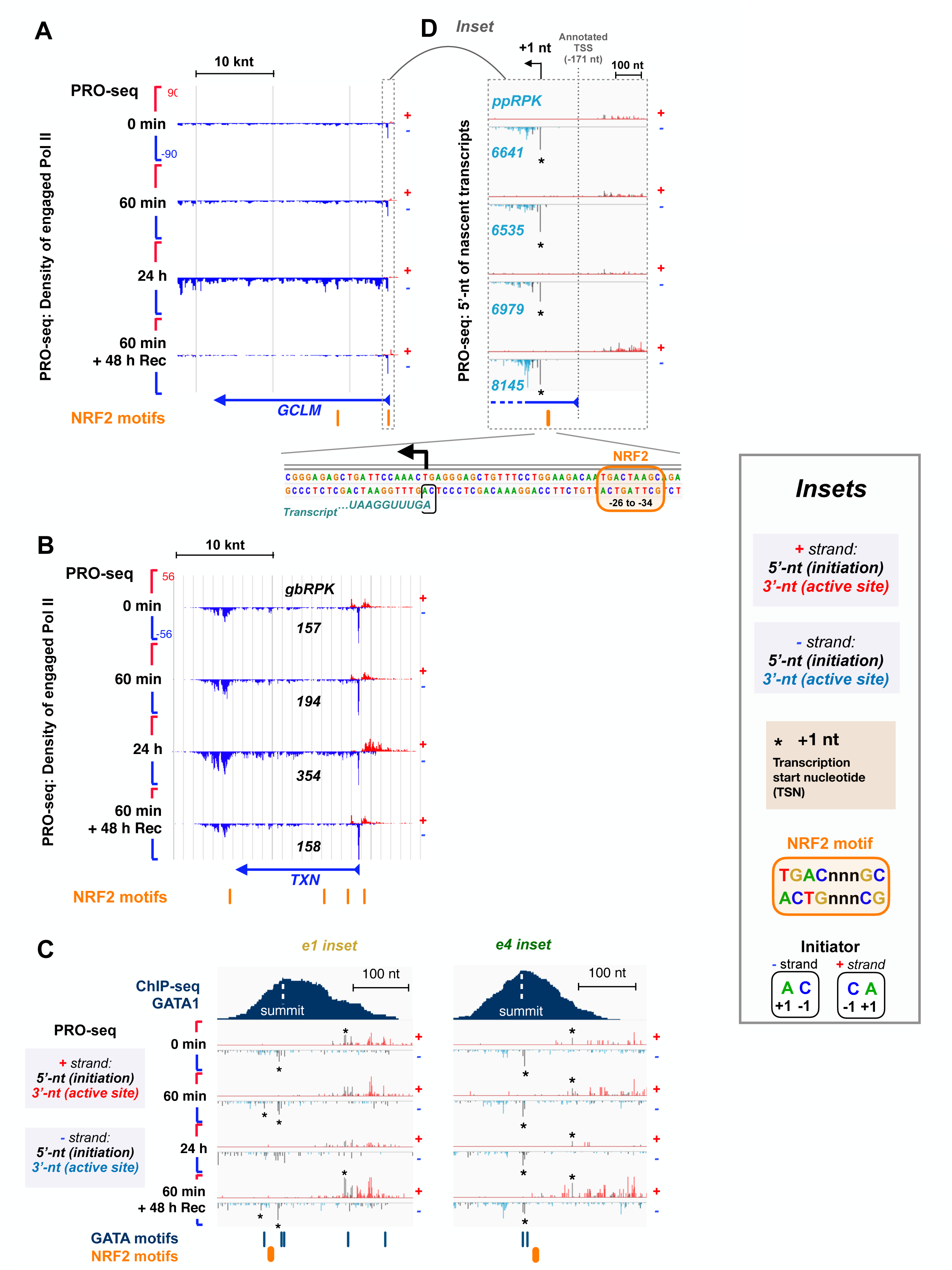
NRF2 directs erythroid differentiation *via* promoters and enhancers. **A-B)** Hemin-induced RNA synthesis at NRF2 target genes **A)** glutamate-cysteine ligase modifier subunit (*GCLM*) and **B)** thioredoxin (*TXN*). **C)** Insets of e1 (left) and e4 (right) of the beta-globin LCR, shown in the main Figure 5D and F. The 5’-ends of nascent transcripts enrich at the precise transcription start nucleotide (+1 nt, TSN) and are indicated with asterisks. The 3’-ends of nascent transcripts (3’nt) show the active site of transcription. **D)** Inset of *GCLM* promoter region, showing the precise transcription start nucleotide (+1 nt, TSN), as identified from the 5’-ends of nascent RNAs (asterisk). NRF2 motif at the promoter of *GCLM* locates -26 nt from the +1 nt.

**Supplementary Figure 6.**
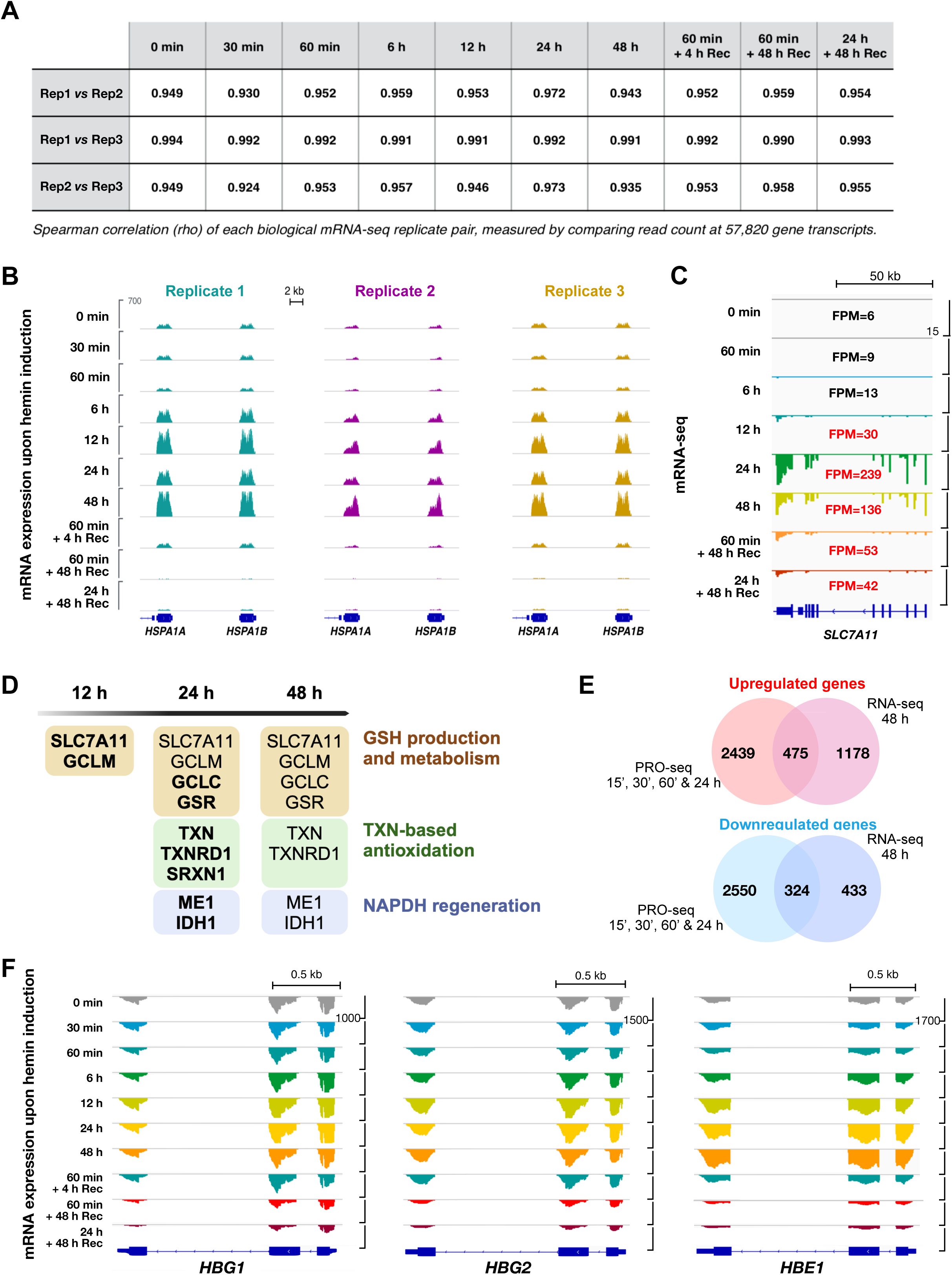
mRNA-seq replicates show high correlation. **A)** Spearman’s rank correlation (rho) of each biological mRNA-seq replicate pair, measured as counts across exons of each gene. **B)** Genome browser showing mRNA levels of *HSPA1A* and *HSPA1B* genes in replicates 1, 2 and 3. **C)** mRNA expression of the NRF2 target gene solute carrier family 7 member 11 (*SLC7A11*). Level of mRNA expression (FPM) is indicated in each condition, red denoting significant induction. **D)** Induced mRNA expression of NRF2 target genes in the glutathione (GSH) metabolism and synthesis (light brown), thioredoxin (TXN) based antioxidation (green) and NADPH regeneration (blue) pathways. The first time point an mRNA is detected induced is shown in bold. **E)** Number of genes associated with changed RNA synthesis within 24 h (PRO-seq), and mRNA expression at 48 h (RNA-seq) of erythroid differentiation. **F)** mRNA expression of the fetal hemoglobin genes *HBG1* and *HBG2*, and the embryonic hemoglobin gene *HBE1* as measured with mRNA-seq upon hemin treatment, and after 4 h or 48 h recovery from a transient hemin exposure.

**Supplementary Figure 7.**
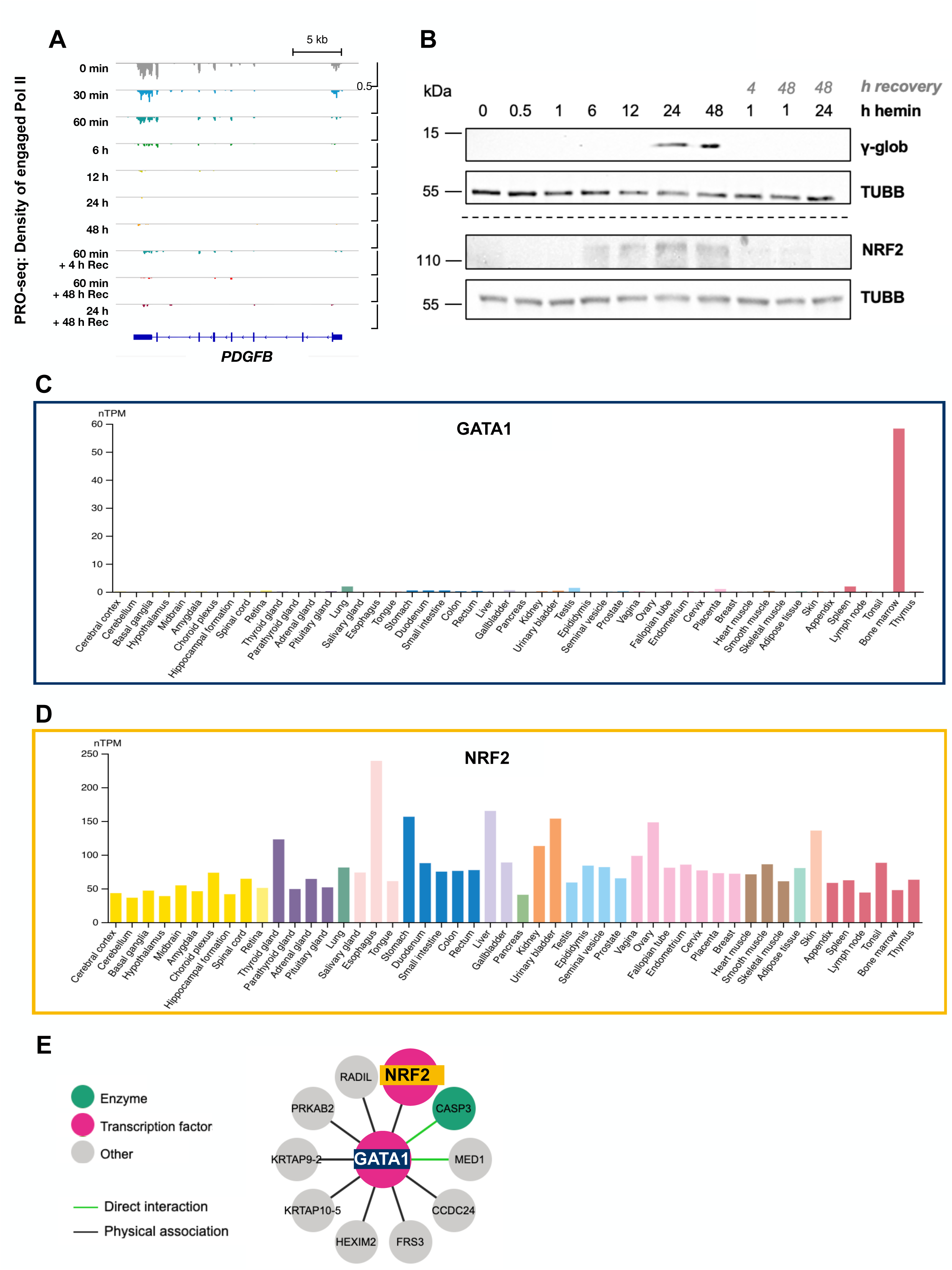
Hemin inhibits megakaryocyte-lineage driving *PDGFB* transcription and increases NRF2 protein levels. **A)** mRNA expression of platelet-derived growth factor subunit B (PDGFB) upon hemin-induced erythroid differentiation. **B)** Immunoblot of γ-globin (γ-glob) and NRF2 protein levels in K562 cells treated with hemin for the indicated time point. For recovery, the hemin was removed, and the cells cultured in hemin-free media. β-tubulin (TUBB) was used as a loading control. **C-D)** Expression on **C)** GATA1 and **D)** NRF2 across human tissues, as identified by the Human Protein Atlas. **E)** GATA1 interactions reported by the Human Protein Atlas. GATA1 and NFE2L2/NRF2 are highlighted. Image credit for C-E: Modified from Human Protein Atlas^51^.

**Supplementary Figure 8.**
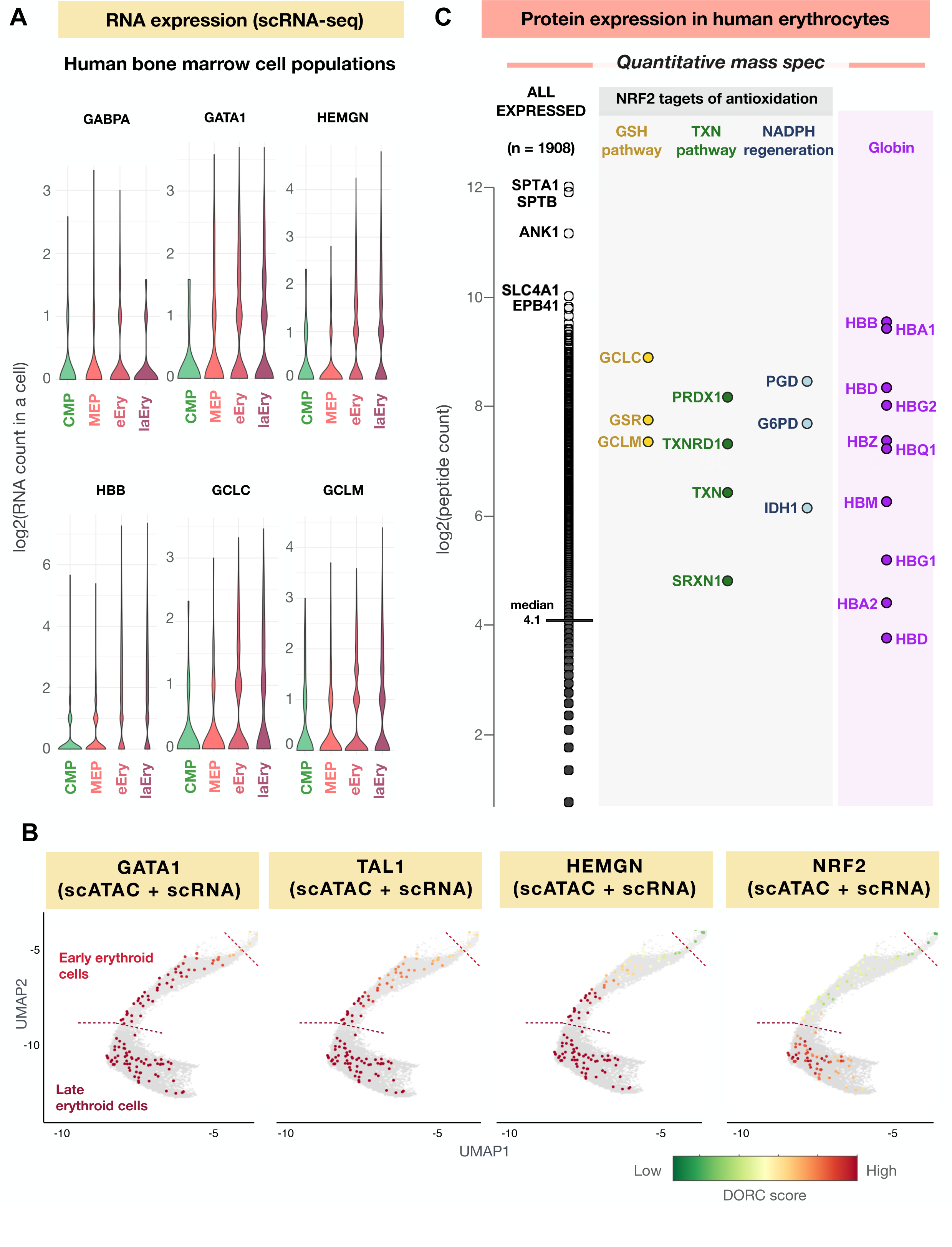
Ordered activation of lineage-specific factors prime for NRF2- driven induction of antioxidation machinery. **A)** Single cell mRNA expression of GABPA, GATA1 and HEMGN, and NRF2 target genes HBB, GCLC and GCLM in indicated human bone marrow cells. The mRNA expression is shown in distinct cell clusters that corresponds to the bone marrow populations shown in the main Figure 6D. **B)** DORC score for GATA1, TAL1, HEMGN and NRF2 in human bone marrow cells, showing their ordered activation during the erythroid lineage specification. The mRNA expression (scRNA-seq) in **A**, and the DORC score (scATAC-seq and scRNA-seq) in **B** originate from SHARE-seq data from the reference^21^. The DORC score in erythroid lineage was visualized in ACAMShiny (https://buenrostrolab.shinyapps.io/ACAMShiny/) and shows the same regions as the main Figure 6D. **C)** Protein expression in human erythrocytes as reported by quantitative mass spectrometry. The peptide counts are shown for all expressed proteins, NRF2 targets genes in the antioxidant pathways, and globins. The quantitative mass spectrometry in panel **C** originates from reference^54^.

